# Proteome profiling of nasopharynx reveals pathophysiological signature of COVID-19 disease severity

**DOI:** 10.1101/2023.07.09.548285

**Authors:** Amanda Ooi, Luke E. Esau, Artyom Pugachev, Arnoud Groen, Sara Mfarrej, Rahul P. Salunke, Amit K. Subudhi, Fathia Ben-Rached, Fadwa Alofi, Afrah Alsomali, Khaled Alquthami, Asim Khogeer, Anwar M. Hashem, Naif Almontashiri, Pierre J. Magistretti, Sharif Hala, Arnab Pain

**Affiliations:** Bioscience Program, Division of Biological and Environmental Sciences and Engineering, 4700 King Abdullah University of Science and Technology, Thuwal 23955-6900, Kingdom of Saudi Arabia; ProteiQ Biosciences GmbH, Am Mühlenberg 11, 14476 Potsdam, Germany; Infectious Diseases Department, King Fahad Hospital, Madinah, Saudi Arabia; Infectious Diseases Department, King Abdullah Medical Complex, Jeddah, Saudi Arabia; Infectious Diseases Medical Department, Al Noor Specialist Hospital, Makkah, MOH, Saudi Arabia; Plan and Research Department, General Directorate of Health Affairs Makkah Region, MOH, Saudi Arabia; Vaccines and Immunotherapy Unit, King Fahd Medical Research Center, King Abdulaziz University, Jeddah, Saudi Arabia; Department of Clinical Microbiology and Immunology, Faculty of Medicine, King Abdulaziz University, Jeddah, Saudi Arabia; College of Applied Medical Sciences, Taibah University, Madinah, Saudi Arabia; Infectious Disease Research Department, King Abdullah International Medical Research Centre, Ministry of National Guard Health Affairs, Jeddah, Saudi Arabia; King Saud bin Abdulaziz University for Health Sciences, Ministry of National Guard Health Affairs, Jeddah, Saudi Arabia; KAUST Smart Health Initiative (KSHI), 4700 King Abdullah University of Science and Technology, Thuwal 23955-6900, Kingdom of Saudi Arabia

**Keywords:** COVID-19 host nasopharynx/ innate immune system/ interferon signaling/ oxidative stress/ retinol metabolism

## Abstract

An aberrant innate immune system caused by the beta coronavirus SARS-CoV-2 is a characteristic manifestation of severe coronavirus disease 2019 (COVID-19). Here, we performed proteome profiling of nasopharyngeal (NP) swabs from 273 hospitalized patients with mild and severe COVID-19 symptoms, including non-survivors. We identified depletion in STAT1-mediated type I interferon response, retinol metabolism and NRF2 antioxidant system that are associated with disease severity in our patient demography. We found that the dysregulation of glucocorticoid signaling and renin-angiotensin-aldosterone system (RAAS) contribute to the pathophysiology of COVID-19 fatality. Hyperactivation of host innate immune system was observed in severe patients, marked by elevated proteins involved in neutrophil degranulation and platelet aggregation. Our study using high-throughput proteomics on the nasopharynx of COVID-19 patients provides additional evidence on the SARS-CoV-2-induced pathophysiological signatures of disease severity and fatality.

## Introduction

COVID-19 caused by severe acute respiratory syndrome coronavirus 2 (SARS-CoV-2), has created an unprecedented global threat, particularly for public health and the global economy due to its considerable mortality and morbidity (Alwan et al., 2020; Blumenthal et al., 2020; Rosenbaum, 2020). There is a dire need to decipher the critical activators and molecular signatures of severe COVID-19, to enable accurate prognosis of future disease courses, and to optimize treatment strategies. Severe COVID-19 can be characterized by an excessive inflammatory response, including a massive cytokine/chemokine expression involving a broad range of immune cells such as macrophages, neutrophils and monocytes. Cytokines are induced downstream of pattern recognition receptors (PRRs), which sense viral molecules (pathogen-associated molecular patterns (PAMPs)) or host molecules associated with disturbance of homeostasis (danger-associated molecular patterns (DAMPs)) (Kawai and Akira, 2010; Mogensen, 2009). Both the induction and function of cytokines, either as antiviral or inflammatory mediators, are highly dependent on the local microenvironment factors (Mehta et al., 2020). SARS-CoV-2-induced viral sepsis releases a number of proinflammatory cytokines such as interleukin (IL)-6, 8 and 17, tumor necrosis factor-*α* (TNF*-α*), and interferon-*γ* (IFN-*γ)*, which correlate with disease severity (Huang et al., 2020). Although inflammation is an essential part of our immune defense to control infection, the inverse correlation between survival and IL-6, IL-8 and TNF-*α* levels (Del Valle et al., 2020; Meizlish et al., 2021) further highlights the potential pathogenic role of an excessive inflammatory response in the pathogenesis of severe COVID-19. A recent analysis of lung immune cells revealed signs of self-sustained inflammatory circuits that might recruit immune cells into the lungs of patients with severe COVID-19 pneumonia (Grant et al., 2021).

Both type I and type III interferon (IFN) responses however, are diminished and delayed in severe COVID-19 patients. This suggests a dual role for type I and III IFN in COVID-19 pathogenesis, with early protection and late amplification of the disease (Galani et al., 2021). SARS-CoV-2 virus has been demonstrated as a poor inducer of IFN-I expression *in vitro*, which is triggered in a delayed manner (Dalskov et al., 2020; Galani et al., 2021; Lei et al., 2020). A transcriptome analysis of peripheral blood mononuclear cells (PBMCs) demonstrates the impaired IFN-∝ production that is often accompanied by a high viral load in severe patients, indicating that IFN-I deficiency could potentially be an immunological hallmark of severe COVID-19 (Hadjadj et al., 2020). The IFN-I (IFN-∝, IFN-*β* and IFN-*ω*) mounts an initial rapid antiviral response as part of the innate immune defense. IFN-I proteins, also a cytokine, are induced when the cell detects viral RNA through Toll-like receptors (TLR3, TLR7 and TLR8) found in endosomes. The IFN-I then binds to the cell-surface receptor IFNAR (IFNAR1 and IFNAR2), resulting in the transcription of IFN-stimulated genes (ISGs) that block the viral replication and transmission (Schoggins et al., 2011). IFN-I deficiency in severe COVID-19 patients might arise through inherited mutations in genes encoding the TLR3 and interferon regulatory factor 7 (IRF7)-dependent IFN-I signaling, rendering the gene products incapable of producing or responding to IFN-I (Zhang et al., 2020c). Besides, inborn genetic defects in TLR7 can predispose to severe COVID-19, providing direct evidence for the importance of IFN signaling in protection against SARS-CoV-2 infection (Asano et al., 2021). A defective antiviral response in severe COVID-19 may also be linked to the development of autoantibodies that bind to and ‘neutralize’ certain IFN-I, thereby inhibiting the antiviral signaling mediated by IFN-∝ and IFN-*ω*. Interestingly, such neutralizing antibodies were mostly found in older males among severe patients (Bastard et al., 2020). Although IFN-I deficiency causes uncontrolled viral replication and infection that can lead to life-threatening COVID-19, this deficiency might also have other immune function consequences, such as the loss of suppression of inflammasomes and enhanced cytokines production downstream of these immune-signaling complexes (Guarda et al., 2011). Such a phenomenon of excessive inflammasome activation might explain severe COVID-19 in IFN-I-deficient individuals.

Retinol depletion and impaired retinoid acid (RA) signaling can contribute to the dysregulation of innate immune system in COVID-19 (Sarohan et al., 2021). Cytosolic viral RNA recognition mechanism through retinoic acid-inducible gene I (RIG-I) receptor (Kawai and Akira, 2008) can rapidly consume a large amount of the host’s retinol reserve, which depletes the serum level of retinol (Chow et al., 2018). The resulting retinol deficiency and impaired RA signaling can impair IFN-I synthesis and excessive inflammatory state in COVID-19 patients (Chattha et al., 2013; Sarohan et al., 2021). Retinol downregulation can disrupt the delicate balance between immune-suppressive regulatory T cells (Treg) and proinflammatory T helper 17 (Th17) cells (Kim, 2008; Raverdeau and Mills, 2014). As such, the coordination between the innate and adaptive components of the immune system is compromised, causing a collapse of the innate immune system and hyperactivation of the Th17 arm of the adaptive immune system, resulting in cytokine storm and systemic organ damage (Casadevall and Pirofski, 2020; Dorward et al., 2021).

The stress-sensing nuclear factor erythroid 2-related factor 2 (NRF2) pathway has been reported to sense the homeostasis alteration during SARS-CoV-2 infection, which shapes the host antiviral and inflammatory responses (Muri and Kopf, 2021). Respiratory viral infections have been associated with inhibition of NRF2-mediated antioxidant pathway and activation of NF-_K_B signaling, which promote oxidative damage and inflammation during infection (Komaravelli and Casola, 2014). In response to oxidative stress, the redox-sensitive NRF2 transcription factor translocate to the nucleus and binds to the cis-acting antioxidant response elements (AREs) to orchestrate the expression of an array of antioxidant genes that regulates the redox homeostasis (Itoh et al., 1999; Raghunath et al., 2018). NRF2 has been shown to mediate the inflammatory response (Thimmulappa et al., 2006) and can transcriptionally repress the inflammatory genes, such as *IL-1β* in murine macrophages (Kobayashi et al., 2016). It has been shown that the NRF2 pathway is suppressed, resulting in oxidative damage in lung biopsies of COVID-19 patients. Importantly, NRF2 agonists can exert potent antiviral action against SARS-CoV-2 infection and block *ex vivo* expression of CXCL10, a pro-inflammatory cytokine, in PBMCs of severe patients (Olagnier et al., 2020). This suggests that SARS-CoV-2 virus impedes the NRF2 pathway to facilitate its replication, while generating a microenvironment prone to pathological hyperinflammation and oxidative stress, contributing to COVID-19 pathogenesis.

Although several omics studies characterizing the comprehensive host response have revealed several molecular signatures of COVID-19 pathology, these investigations are mostly performed in the patient’s serum, urine, tissue biopsies, and to a small extent, in the NP swab. In this study, we captured the SARS-CoV-2 induced molecular phenotype in the nasopharynx of 273 hospitalized patients with mild and severe symptoms, including non-survivors, by measuring the nasopharyngeal proteome in an untargeted fashion. To achieve this, we applied the use of data-independent acquisition mass spectrometry (DIA-MS) coupled with *infineQ* data processing software, utilizing the advantage of high-flow chromatography in short-gradient proteomics to enable high quantification precision, improved data acquisition and processing workflow. Samples were collected during the first wave of early COVID-19 pandemic, between March and August 2020. Here, we show the hyperactivation of host innate immune system drives the heightened inflammatory response and oxidative stress, as evidenced by elevated neutrophil degranulation, interleukin signaling, and platelet activation in severe patients. In parallel, we observed the decreased STAT1-mediated IFN-I response, retinol metabolism, and NRF2 antioxidant system that are associated with disease severity. We also found that dysregulation in glucocorticoid signaling and RAAS can contribute to the pathophysiology of COVID-19 fatality. As such, our study provides evidence of specific molecular pathways and mechanisms underlying the pathophysiology of severe COVID-19 that can be associated with the aberrant innate immune system caused by the SARS-CoV-2 virus.

## Results & Discussion

### A high-throughput DIA-based quantitative proteome profiling of COVID-19 NP swabs

We collected NP swabs from a cohort of 273 hospitalized COVID-19 patients across the region of Jeddah, Makkah, and Madinah in Saudi Arabia during the early pandemic between March to August 2020. Our patients’ cohort consists of 92 mild (M), 91 severe alive (SA), and 90 severe fatal (SF) cases respectively (**Figure 1A**). We classify the maximal severity of COVID-19 retrospectively by determining the presence of symptoms, the need for oxygen supplementation and the level of respiratory support. M patients are denoted as symptomatic without oxygen requirement or respiratory support, whereas both SA and SF groups have received either respiratory support or mechanical ventilation. In total, there are 207 male and 66 female COVID-19 patients in three age categories ranging from 25 – 45 (n = 79), 46 – 55 (n = 88) and 56-and-above (n = 106) years old recruited for this study (**Figure 1A**, **Table 1**). This study serves to understand the pathogenesis of COVID-19 disease severity and fatality among hospitalized COVID-19 patients by performing a DIA-based quantitative proteome profiling of the host nasopharynx.

**Figure 1.**
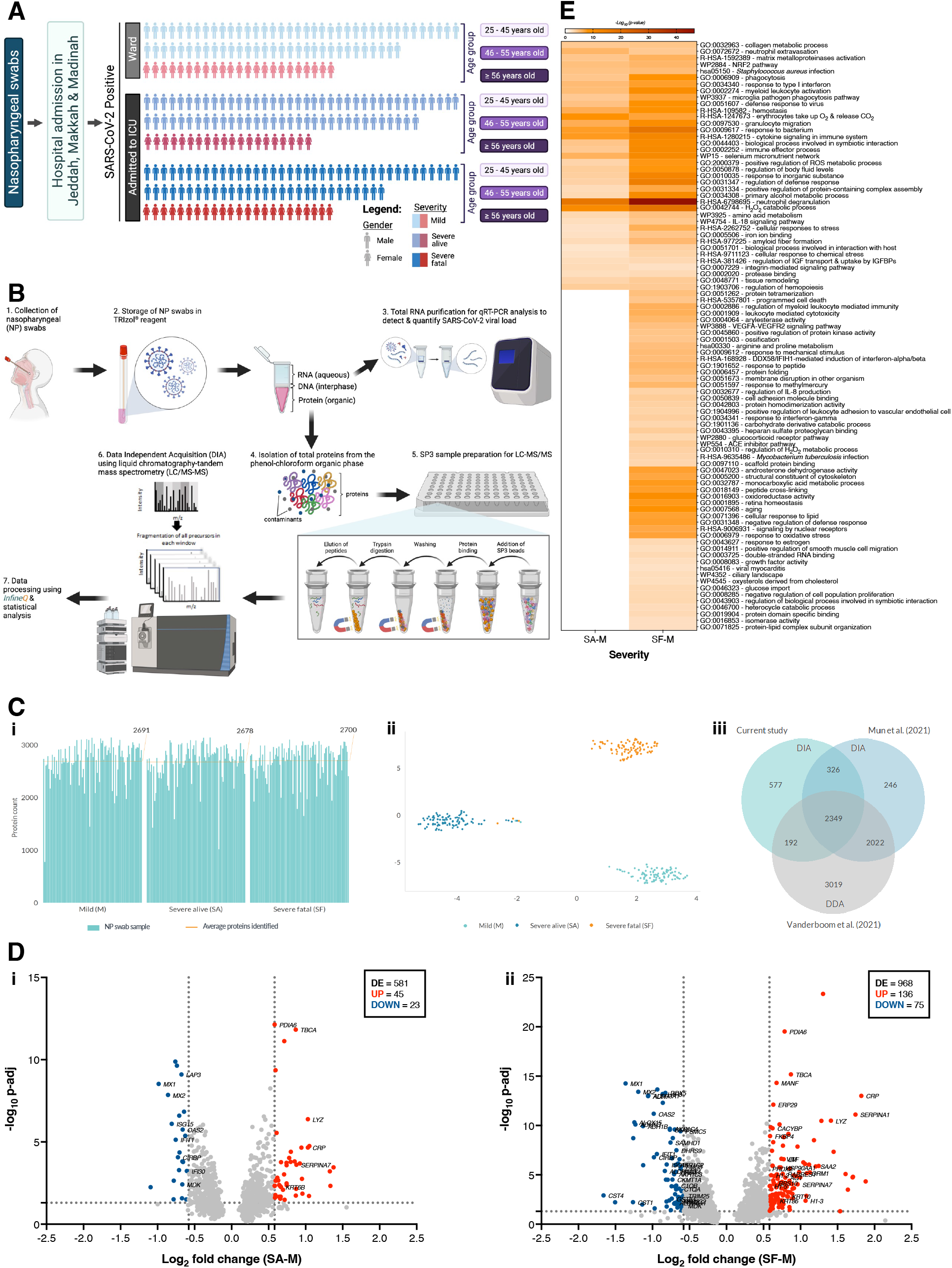
Comparative proteome profiling of COVID-19 patients NP swabs using a DIA-based LC-MS analysis. **A)** The demography of our enrolled COVID-19 patients. **B)** A workflow showing the collection of NP swab in TRIzol^®^ reagent followed by total RNA and protein extraction from the aqueous and organic phase respectively. The RNA was analyzed by qRT-PCR to measure the SARS-CoV-2 viral load prior to protein sample preparation using an adapted SP3 method for a DIA-based LC-MS/MS analysis. The raw data was then processed using *infineQ* followed by statistical analyses. **C)** (**i**) Identification of protein group number for each sample. The orange line drawn across the bar chart represents the average number of identified protein groups for each severity, and the average protein number is indicated at the top right of each group. (**ii**) Cluster forming along two linear dimensions of LDA analysis based on principal components explaining 95% variance (PC-LDA) showing almost perfect separability of classes. (**iii**) A comparison of our proteome data with other published COVID-19 nasopharynx proteomes that used DIA (Mun et al., 2021) and DDA(Vanderboom et al., 2021) approaches. **D)** The differential protein expression relating to COVID-19 severity for (**i**) severe alive vs mild (SA-M) and (**ii**) severe fatal vs mild (SF-M) respectively. Using a robust linear mixed effect modeling, the total number of differentially expressed (DE) proteins identified were 581 and 968 for SA-M and SF-M respectively, after FDR correction with an adjusted p-value cut-off of 5 % (p-adj < 0.05) performed using a Benjamin-Hochberg analysis. Red and blue circles highlight the upregulated (UP) and downregulated (DOWN) proteins respectively with an average fold change of ≥ 1.5 (-0.58 ≤ log_2_ fold change ≥ 0.58) and a p-adj < 0.05 (-log_10_ p-adj > 1.3). **E)** Top 90 enriched terms for DE proteins in SA-M and SF-M groups, as represented by p-values and analyzed using Metascape (version 3.5.20211101). The p-value significance is represented by a color shade ranging from lighter orange (less significant) to red (more significant), while white color indicates a lack of significance.

**Table 1.** Demographics and baseline characteristics of COVID-19 patients.

| Variables | COVID-19 |  |  |
| --- | --- | --- | --- |
|  | MILD | Severe Alive | Severe Dead |
| **Gender - no. (%)** |  |  |  |
| Male | 69 (75%) | 71 (78%) | 67 (74.4%) |
| Female | 23 (25%) | 20 (22%) | 23 (25.6%) |
| **Age - year** |  |  |  |
| Mean ± SD | 51.06 ± 11.9 | 51.1 ± 12.12 | 55.4 ± 12.6 |
| Range | 23 - 82 | 23 - 82 | 25 - 82 |
| **Region** |  |  |  |
| Middle East (ME) | 48 | 55 | 51 |
| South Asia (SA) | 42 | 34 | 35 |
| South East Asia (SEA) | 1 | 0 | 4 |
| Other | 1 | 2 | 0 |
| **Comorbidity** |  |  |  |
| Overweight | 1 | 3 | 1 |
| Cardiovascular | 4 | 9 | 10 |
| Hypertension | 36 | 37 | 35 |
| Diabetes | 33 | 34 | 34 |
| **Treatment - no. (%)** |  |  |  |
| Oxygen | 0 (0%) | 2 (2.2%) | 1 (1.1%) |
| Mechanical ventilation | 0 (0%) | 37 (40.6%) | 87 (96.7%) |
| **Other** |  |  |  |
| Viral CT (Mean ± SD) | 30.3 ± 5.01 | 31.2 ± 5.13 | 27.9 ± 5.16 |

**Table 2.**
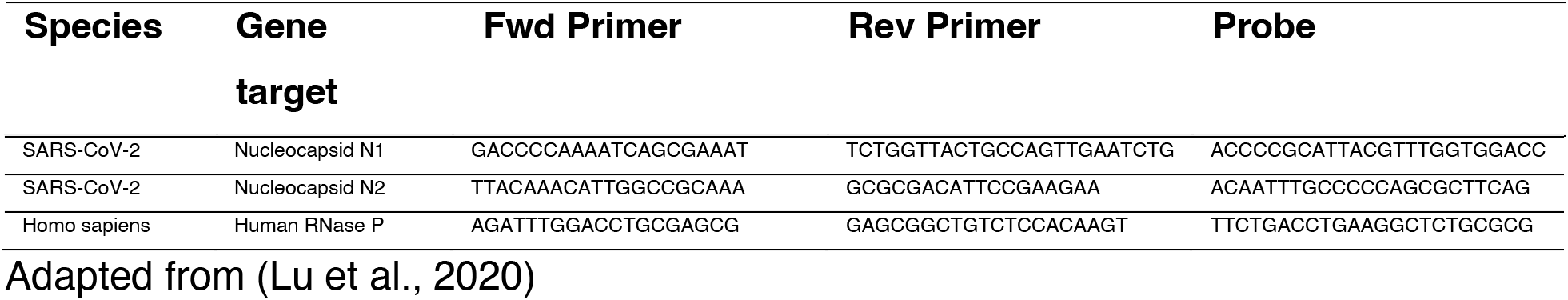
Primer/probe sequences used in this study for the detection of SARS-CoV-2 via qRT-PCR.

**Table 3.**
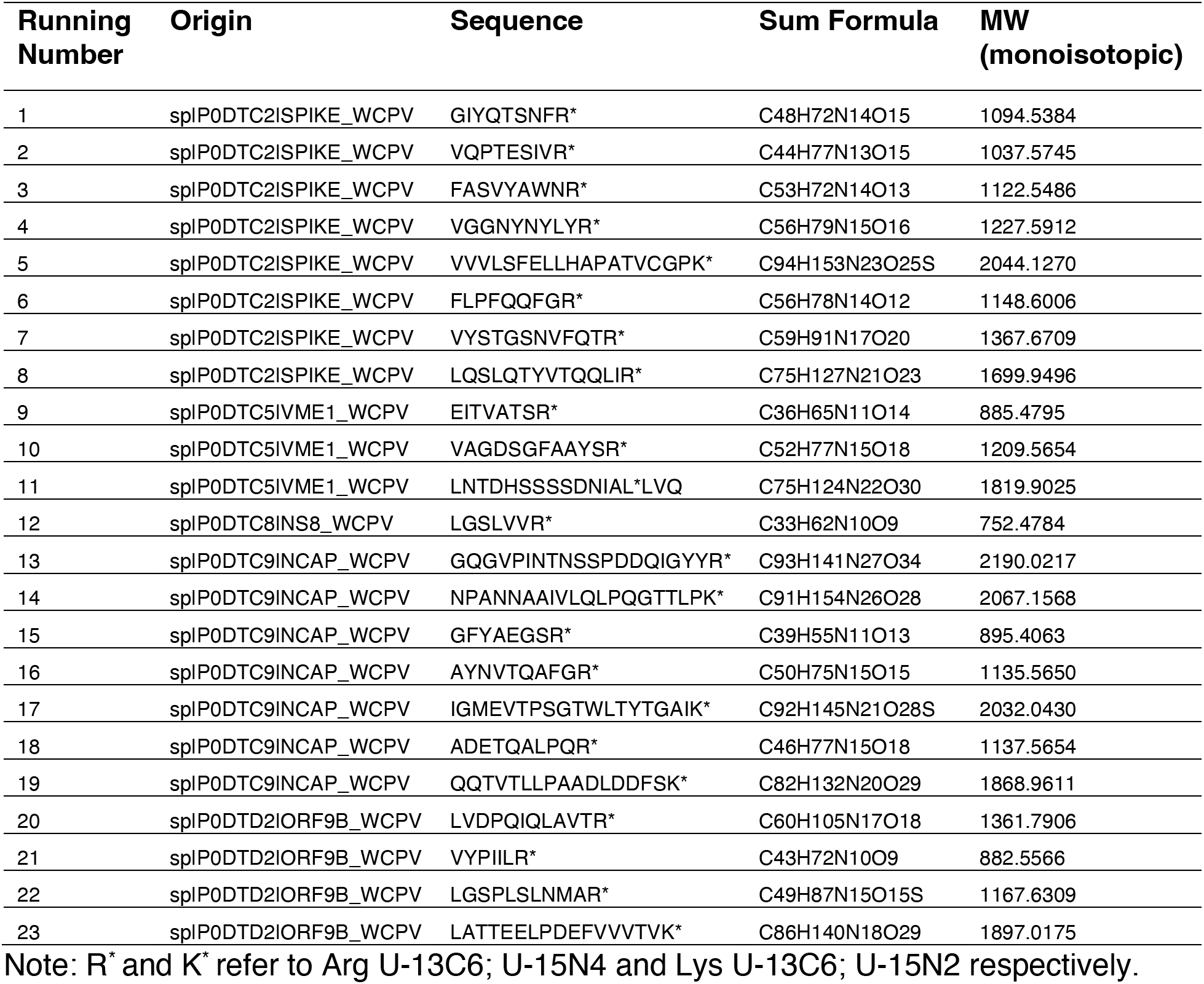
Peptide sequences of the SpikeMix^TM^ SARS-CoV-2-heavy.

Here, we developed a high-throughput workflow (**Figure 1B**) from sample preparation to peptide measurement by liquid-chromatography tandem mass spectrometry (LC-MS/MS), and finally to data processing and analysis using *infineQ* software. Briefly, all NP swabs were collected and stored in TRIzol^®^ reagent after which, we extracted both the total RNA and proteins from each sample. We then performed qRT-PCR on the total RNA to measure the SARS-CoV-2 viral load (a median Cycle threshold (Ct) value of ≤ 37, i.e. greater than four viral copies were selected for this study), ensuring that all samples included in this study are COVID-19 positive prior to peptide preparation for LC-MS/MS analysis. In order to process large number of protein samples, we modified the single-pot, solid-phase-enhanced sample-preparation (SP3) method (Hughes et al., 2019) to perform the entire peptide digestion and clean-up process in a 96-well plate format. The SP3 workflow adopts the paramagnetic bead-based approach that uses hydrophilic interaction mechanism and ethanol-driven solvation to efficiently clean up and recover proteins/peptides in a single tube (Hughes et al., 2019). We applied a DIA-based MS data acquisition approach (**Figure S1**) to measure the peptides of individual clinical NP swab. In combination with a specific spectral library, we identified an average of 3545 protein groups per sample (**Figure 1Ci**) with a technical coefficient variation (CV) of 21 % of pooled protein samples (**Figure S1Bii**). The biological variation from the clinical samples allowed nearly perfect patient class separation using PC-LDA analysis (**Figure 1Cii**).

We then assessed the comprehensiveness of our proteome by comparing it to previous proteome studies on COVID-19 NP swabs (**Figure 1Ciii**). Overall, we identified fewer protein groups in all 273 samples. (Mun et al., 2021) detected a total of 4943 protein groups in 90 NP swabs (45 COVID-19 negative and 45 COVID-19 positive) using a DIA-PASEF approach, whereas (Vanderboom et al., 2021) identified 7582 proteins in 16 NP samples (8 COVID-19 negative and 8 COVID-19 positive) using extensive sample pre-fractionation prior to DDA-TMT MS data acquisition. We observed an overlap of 2675 protein groups between our study with (Mun et al., 2021) which used a similar approach, while 2349 proteins were shared across all three studies (**Figure 1Ciii**). As our sample size is much larger, we can arguably confirm that our study has provided a protein catalog that is of high reproducibility from COVID-19 positive NP swabs. NP swabs contain a heterogenous mixture of immune (dendritic and macrophages) and epithelial (ciliated and goblet) cells (Poulos et al., 2020). To verify our samples, we performed a tissue or cell specific enrichment analysis (**Figure S2A**) using the PaGenBase database (Pan et al., 2013). We found that our NP samples are enriched with bronchial epithelial cells, and immune cell subtypes including macrophages, CD33+, CD34+, and CD71+-expressing immune cells (**Figure S2A**). This is consistent with the cell subtype analysis performed by (Mun et al., 2021) using Enrichr (Kuleshov et al., 2016), in which we identified similar cell subtypes, except for B lymphoblasts, dendritic and natural killer cells.

Further analysis using a robust linear mixed effect modeling was performed on the 3545 identified proteins, followed by FDR correction using the Benjamin-Hochberg method with an adjusted p-value cut-off of 5%. After correction, we identified 1236 differentially expressed (DE) proteins across all samples. Differential analyses between severe alive and mild patients (SA-M) (**Figure 1Di**) and between deceased severe and mild patients (SF-M) (**Figure 1Dii**) reveal a more pronounced number of DE proteins in SF (968) than in SA (581) group (adjusted p-value of 0.05). Among the DE proteins in SF-M and SA-M groups respectively, 136 (SF-M) and 45 (SA-M) proteins were upregulated, while 75 (SF-M) and 23 (SA-M) proteins were downregulated (≥ 1.5 fold-change) (**Figure 1D**). Using the lists of DE proteins (≥ 1.5 fold-change) in SA-M and SF-M, we performed a comparative analysis of these datasets on Metascape (Zhou et al., 2019). Overall, the functional enrichment analyses across these DE proteins show that the alteration of host proteome in SF is more perturbed than in SA patients (**Figure 1E**). A cross comparison between SA-M and SF-M DE proteins also reveals that 75.7 % of DE proteins in SA overlapped with SF proteins, whereas, half of the DE proteins (51.5 %) is unique to SF group itself (**Figure S2B**).

### IFN-I deficiency in severe COVID-19 via aberrant STAT1 signaling enables SARS-CoV-2 virus to evade the host antiviral response for its replication

Functional enrichment analysis on significantly downregulated proteins using the PANTHER classification system (Mi et al., 2021) shows that IFN signaling is impaired in severe patients (**Figure 2A** and **Figure S3A**). To further understand the effect of SARS-CoV-2 infection on IFN signaling in the nasopharynx of severe patients, we performed a comparative search on the downregulated proteins using Interferome (Rusinova et al., 2013). Nearly half (47.8 %) of the downregulated proteins (11 out of 23 proteins) in SA patients were identified as IFN-regulated genes (IRGs), whereas the IRGs comprise 20 % of the downregulated proteins (15 out of 75 proteins) in SF group (**Figure 2Bi**). While most of the identified IRGs proteins belong to IFN-I, nine proteins (CIRBP, DGKZ, IFIT1, ISG15, LAP3, MDK, MX1, MX2, and OAS2) are overlapped between the SA and SF groups (**Figure 2Bi**). Central to antiviral immunity, these nine IFN-I proteins are classical ISGs involved in various antiviral functions against SARS-CoV, Middle-East Respiratory Syndrome (MERS), SARS-CoV-2 and other respiratory syncytial viral infections (Blanco-Melo et al., 2020; Hadjadj et al., 2020; King and Sprent, 2021; Masood et al., 2021; Schneider et al., 2014; Vanderboom et al., 2021). Although IFN-I plays a crucial role as the first line of defense against viruses, there is still a discrepancy regarding its role in COVID-19 (King and Sprent, 2021; Lee and Shin, 2020; Zanoni, 2021). Robust IFN-I responses have been reported, associating the increase in IFN-I response with immunological characteristic of patients with severe COVID-19 (Lee and Shin, 2020; Zanoni, 2021). On the other hand, a PBMCs transcriptome study shows the impaired IFN-∝ production and activity is often accompanied by a high viral load in severe patients, suggesting that IFN-I deficiency could potentially be an immunological hallmark of severe COVID-19 (Hadjadj et al., 2020). Similarly, a downregulation of ISGs in the blood of severe COVID-19 patients was observed (Masood et al., 2021). A single-cell RNA sequencing (scRNA-seq) analysis on all major immune blood cell types revealed that severe COVID-19 patients exhibit a lack of ISGs-expressing cells as compared to mild individuals. A lack of coordinated pattern of ISGs expression across every blood immune cell population in severe patients was associated with high levels of antibodies that functionally inhibit the production of ISG-expressing cells by antagonizing the initial IFNAR signaling via Fc *γ* receptor IIb (Fc*γ*RIIb) (Combes et al., 2021).

**Figure 2.**
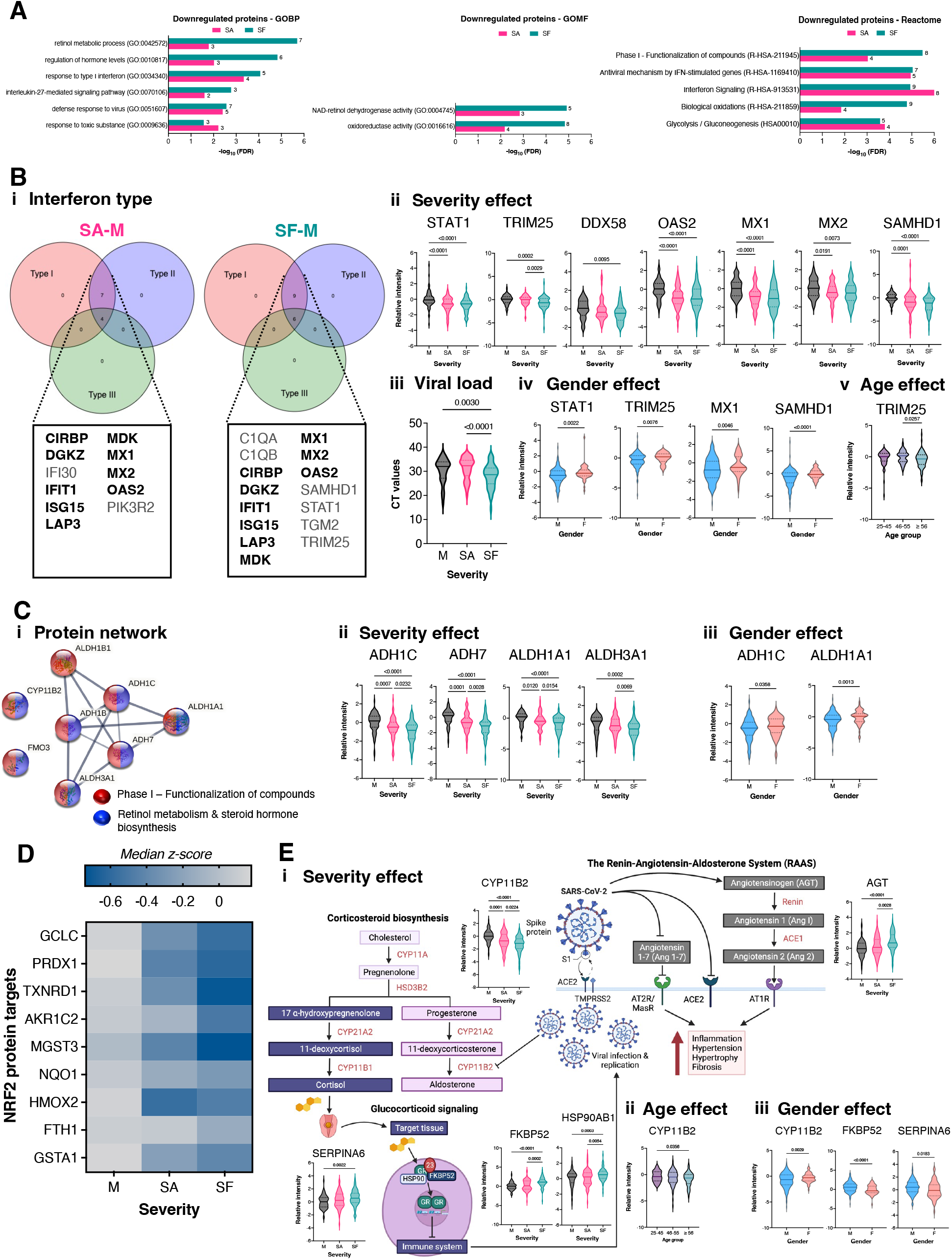
SARS-CoV-2 virus impedes the STAT1-mediated IFN-I production, retinol metabolism, NRF2 antioxidant, glucocorticoid, and RAAS signaling pathways. **A)** Gene ontology (GO) enrichment analysis of significantly downregulated proteins (≥ 1.5 fold-change) in SA and SF groups. Using the PANTHER classification system, a statistical over-representation test on the list of downregulated proteins was performed with Fisher’s exact test with FDR multiple test correction, set at a critical value of FDR < 0.05. The protein function is classified based on complete Gene Ontology Biological Process (GOBP) and Gene Ontology Molecular Function (GOMF), while the pathway analysis is mapped using the Reactome database. The number of proteins for each GO/Reactome term is annotated at the end of respective bar graphs. **B)** Mapping of downregulated proteins onto Interferome database (version 2.01) has identified 11 and 15 IFN-regulated proteins for SA-M and SF-M group respectively. (**i**) Majority of downregulated IFN proteins belong to type I interferon (IFN-I). The overlapped IFN-regulated proteins between SA-M and SF-M are listed in bold. (**ii**) Differential protein expression level for selected IFN-I-regulated proteins based on *z*-scores calculated against the median of mild patients. (**iii**) CT values representing the SARS-CoV-2 viral load. (**iv**) Differential protein expression levels of STAT1, TRIM25, MX1 and SAMHD1 between male (M) and female (F). (**v**) Age effect on TRIM25 protein expression. Patients are grouped into three age categories, ranging from 25 to 45, 46 to 55 and 56 years-old and above. **C)** (**i**) Protein network analysis of downregulated proteins involved in phase 1 of drug metabolism (red nodule) and retinol metabolism (blue nodule) using STRING version 11.5. The network nodes represent the downregulated proteins, while the line thickness indicates the degree of confidence prediction of functional association. (**ii** – **iii**) Median-normalized protein abundance of ADH1C, ADH7, ALDH1A1, and ALDH3A1versus disease severity (**ii**) and gender (**iii**) respectively. **D)** Downregulation of a repertoire of NRF2-mediated ARE-dependent antioxidant proteins versus COVID-19 severity. Protein abundances were plotted based on the median *z*-score, ranging from blue (low expression) to gray (high expression). **E)** (**i – iii**) Pathways of corticosteroids biosynthesis from cholesterol, glucocorticoid signaling and the renin-angiotensin-aldosterone system (RAAS) with selected differentially expressed proteins indicate a certain degree of dysregulation in these pathways with respect to disease severity (**i**), age (**ii**) and gender (**iii**). Data information: *p*-values for all violin plots were calculated based on ordinary one-way ANOVA followed by Tukey’s multiple comparisons test for COVID-19 patient’s severity (n_M_ = 92; n_SA_ = 91; n_SF_ = 90) and age groups (n_25-45_ = 79; n_46-55_ = 88; n_≥56_ = 106), whereas the gender factor (n_M_ = 207; n_F_ = 66) was analyzed using the unpaired t-test with Welch’s correction.

Here, we show that OAS2, MX1 and MX2 are downregulated in severe patients (**Figure 2Bii**). Together with RNase L, OAS2 plays an important role in early viral clearance through viral RNA degradation (Masood et al., 2021). MX1 and MX2 differ considerably in their viral specificities and mechanisms of action, with MX1 exhibiting a wider antiviral action against various RNA and DNA viruses (Haller et al., 2015), while the antiviral activity of MX2 pertains to certain viruses such as HIV (Bhargava et al., 2018). Specifically acting on the viral ribonucleoprotein complex, MX1 has been identified as a critical responder in SARS-CoV-2 infection that could potentially be a therapeutic target for COVID-19 (Bizzotto et al., 2020). Consistent with this notion, a genome-wide association study on COVID-19 patients with European genetic ancestry shows that the minor alleles of five single nucleotide polymorphisms within *TMPRSS2* and near *MX1* gene correlate with a reduced risk of developing severe COVID-19 (Andolfo et al., 2021). Together, both studies support the evidence that MX1 acts as an antiviral effector against SARS-CoV-2, thereby determining the COVID-19 disease progression (Andolfo et al., 2021; Bizzotto et al., 2020). Indeed, we found that the MX1 protein expression negatively correlates with the disease severity (**Figure 2Bii**) and its downregulation is associated with increased viral load (**Figure 2Biii**). We also observed a gender bias in MX1 expression. Females tend to show a higher protein level of MX1, suggesting that females mount a stronger IFN-driven antiviral response than males (**Figure 2Biv**). It has been reported that COVID-19 female patients have higher plasma concentrations of IFN-*α* over the disease course (Takahashi et al., 2020), while autoantibodies that inhibit IFN-I signaling have been reported in a subset of severe COVID-19 patients, majority (94 %) of whom were older males (Bastard et al., 2020).

STAT1, TRIM25, DDX58, and SAMHD1 that are involved in IFN-I signaling are also downregulated in severe patients (**Figure 2Bii** and **Figure S3A**). DDX58/ retinoic acid-inducible gene I protein (RIG-I) is an antiviral innate immune response receptor that senses viral 5’ tri-phosphorylated RNA and short double-stranded RNA species, and induces a downstream IFN signaling cascade (Pichlmair et al., 2006). DDX58 does not only potently activates the IFN expression, but can also induce the production of proinflammatory cytokines (Mogensen, 2009; Paludan and Mogensen, 2022). Prior to activation of DDX58-mediated IFN signaling through an ubiquitination-dependent mechanism, DDX58 forms a ribonucleoprotein complex with viral RNAs on which it homo-oligomerizes to form filaments in order to recruit the E3 ubiquitin/ISG15 ligase (TRIM25). TRIM25 induces the Lys 63-linked polyubiquitination of DDX58 to activate and facilitate the interaction of DDX58 with mitochondria antiviral signaling protein (MAVS), which is crucial for the DDX58-mediated interferon-*β* production and antiviral activity (Gack et al., 2007). Recently, it has been shown that the SARS-CoV-2 viral nucleocapsid protein (N protein) binds to the SPRY domain of TRIM25, and interferes with the TRIM25-DDX58 interaction. This in turn impedes the TRIM25-mediated DDX58 ubiquitination and activation, thereby inhibits the IFN-*β* production (Gori Savellini et al., 2021). We found that the expression levels of DDX58 and TRIM25 are suppressed by SARS-CoV-2, with a stronger inhibitory effect observed in SF patients (**Figure 2Bii**). We also observed a significant gender difference in the expression of TRIM25, but not in DDX58 (**Figure 2Biv**), whereby the TRIM25 protein level is higher in females. The expression of TRIM25 is also significantly depleted in older patients (≥ 56 years old) (**Figure 2Bv**). Taken together, our finding corroborates that the SARS-CoV-2 virus hijacks the innate immune signaling in severe patients by: 1) directly interfering with the TRIM25-mediated DDX58 activation, and 2) impairing the ISGs expression in the downstream effector machinery of IFN, as evidenced by the depletion of IFN-I antiviral response (**Figure 2A-B**) to circumvent the host defense.

We found that SARS-CoV-2 also has the ability to impair the signal transducer and activator of transcription 1-alpha/beta (STAT1)-mediating cellular responses to IFNs and other cytokines family. STAT1 was significantly downregulated in severe patients (**Figure 2Bii**). This observation is consistent with a recent report showing a depletion in STAT1 expression in CD14^+^ monocytes and plasma blasts of severe COVID-19 patients (Rincon-Arevalo et al., 2022). We noticed that the STAT1 protein expression is significantly lower in males (**Figure 2Biv**). It has been demonstrated that the SARS-CoV-2 N protein binds to and inhibits both the phosphorylation and nuclear translocation of STAT1 and STAT2, thereby antagonizing the IFN-I signaling in order to replicate efficiently in the host cells (Mu et al., 2020). The SARS-CoV-2-mediated inhibition of STAT1 phosphorylation has been shown to attenuate the ISGs transcription in monocyte-derived dendritic cells and macrophages (Yang et al., 2020). These findings suggest that an aberrant STAT pathway could be a crucial determinant for severe COVID-19 pathogenesis. Several mechanisms have been proposed to explain how a SARS-CoV-2-driven dysfunction in STAT1 could lead to compensatory hyperactivation of STAT3 that could further inhibit the STAT1-mediated IFN-I response. The aberrant transcriptional rewiring, starting with the downregulation of STAT1 and shifting towards STAT3 hyperactivation may lead to severe COVID-19 symptoms such as coagulopathy, thrombosis, increase in inflammation, profibrosis, and T cell lymphopenia (Matsuyama et al., 2020).

Furthermore, we show that the sterile alpha motif and HD-domain-containing protein 1 (SAMHD1) is significantly downregulated in severe patients (**Figure 2Bii**). The depletion of SAMHD1 protein is associated with increased disease severity with a concomitant increase in viral load, particularly in SF patients (**Figure 2Bii-iii**). We also found a significantly higher expression level of SAMHD1 in females (**Figure 2Biv**). Overall, we identified an impaired IFN-I response against SARS-CoV-2 infection in severe patients that is accompanied by a high viral load in the nasopharynx, the SF group in particular (**Figure 2Biii**). We also found an association of IFN-I deficiency with gender and age in COVID-19 (**Figure 2Biv-v**). We observed that the activation of antiviral response is differentially regulated between men and women, and can be inversely correlated with age. In general, women develop stronger innate and adaptive immune responses against most viral infections (Bunders and Altfeld, 2020; Klein and Flanagan, 2016). We found a significant sex difference in the induction of IFN-I response that is critical for restriction of SARS-CoV-2 viral replication (**Figure 2Biv**). However, we are aware that there is a potential bias in interpreting this result due to the significantly fewer females included in our patient cohort.

### Dysregulation in the retinol metabolism, NRF2, glucocorticoid, and RAAS signaling pathways contribute to the pathophysiology of COVID-19 severity and fatality

Early reports initially attributed the profound defect in IFN-I production in severe COVID-19 to mutations in genes involved in regulating type I and III IFN immunity (Zhang et al., 2020c). However, the aberrant genes encoding TLR3- and IRF7-dependent IFN-I immunity at eight loci were only found in 3.5% (23 out of 659 severe COVID-19 patients) of their cohort. These mutations alone could not fully explain the impairment of IFN-I production in severe COVID-19 (Zhang et al., 2020c). Such mutations bearing a critical role in the host innate immune system could have been lethal early in life due to exposure to various infections. Therefore, we hypothesize that the depletion in retinol metabolism followed by downregulation of RA signaling (**Figure 2C**) may contribute to the deficiency in IFN-I production. Among the seven downregulated proteins involved in retinol metabolism and steroid hormone biosynthesis (**Figure 2Ci**), we show for the first time a pronounced effect in the downregulation of ADH1C, ADH7, ALDH1A1, and ALDH3A1 proteins, which negatively correlates with COVID-19 severity (**Figure 2Cii**). ADH1C and ALDH1A1 proteins are differentially regulated between genders, with higher protein levels observed in females (**Figure 2Ciii**). As SARS-CoV-2 virus has a large genome size of 30 kB (Harrison et al., 2020; Naqvi et al., 2020), it can release large amounts of RNA particles in the host cells. This leads to prolonged and overwhelming immune stimulation, and the increased RA consumption over time depletes the host retinol reservoir. As such, the DDX58-mediated IFN signaling collapses (**Figure 2Bii**) and disrupts the fine balance between immune-suppressive Treg cells and proinflammatory Th17 cells in severe COVID-19 (Kim, 2008; Raverdeau and Mills, 2014). The lack of RA signaling in severe patients fails to block the IL-6 signaling, leading to the expression and activation of RA-related orphan receptor gamma (ROR *γt*) transcription factor that promotes the differentiation of naïve T cells into proinflammatory Th17 cells (Chen et al., 2011; Elias et al., 2008; Lu et al., 2014). As a result, excessive cytokine discharge due to hyperactivation of NF-_K_B signaling in Th17 cells generates a hyperinflammatory response in severe COVID-19. It has been shown that the level of anti-inflammatory Treg cells in severe COVID-19 is lower than the mild patients, which may further contribute to uncontrolled immune reactions (Tan et al., 2020).

Here, we show that RA synthesis is suppressed in severe patients, as evidenced by the downregulation of ADH7 and ALDH1A1. ADH7, a NAD^+^-dependent all-trans-retinol dehydrogenase, is involved in the oxidation of all trans-retinol into retinaldehyde, and ALDH1A1 functions downstream of retinol dehydrogenases, catalyzes the oxidation of retinaldehyde into RA, which is the second step in the metabolism of retinol into RA (**Figure 2Cii**). This suggests that the decrease in RA synthesis and signaling caused by SARS-CoV-2 infection disrupts the production of IFN-I in severe COVID-19, thereby enabling the replication, transmission and pathogenesis of SARS-CoV-2 without any hindrance in the host cells (**Figure 2A-C**). Our data corroborate a recent finding showing the depletion in serum retinol levels in severe COVID-19 patients that is independent of age and comorbidity (Sarohan et al., 2022). Another study has demonstrated a potent antiviral action of all-trans retinoic acid (ATRA) against SARS-CoV-2 virus by interrupting the spike-mediated cellular entry in both human cell lines and organoids of the lower respiratory tract (Tong et al., 2022). It has also been shown that retinol metabolism and RA signaling can function as a critical host-pathogen-interaction circuit in controlling SARS-CoV-2 infection. The RA receptor (RAR) agonists can impede viral replication, in which the inhibition was reversed with RAR antagonists, implicating the importance of RAR-mediated signaling in the pathogenesis of COVID-19 (Riva et al., 2020). As retinol is the biological active form of vitamin A, vitamin A and its derivatives strengthen the mucosal immunity and barrier formation against viral infection by increasing the production of IFN-I and secretory IgA (Le Page et al., 2000; Onoguchi et al., 2011; Trasino, 2020). A significant depletion in plasma level of vitamin A was found in severe COVID-19. Patients with < 2 mg/L of plasma vitamin A level were found to be significantly associated with the development of ARDS and mortality. The reduced level of vitamin A correlates significantly with increased levels of inflammatory markers including C-reactive protein (CRP) and ferritin, and incidence of lymphopenia in severe patients (Tepasse et al., 2021). In parallel, we also observed significantly enhanced CRP expression in the nasopharynx of severe patients, which is higher in SF (**Figure 3Di** and **Figure S3Gi**). Therefore, reconstitution of RA signaling with vitamin A supplementation may provide a valuable strategy for management of COVID-19, which requires further clinical investigations.

**Figure 3.**
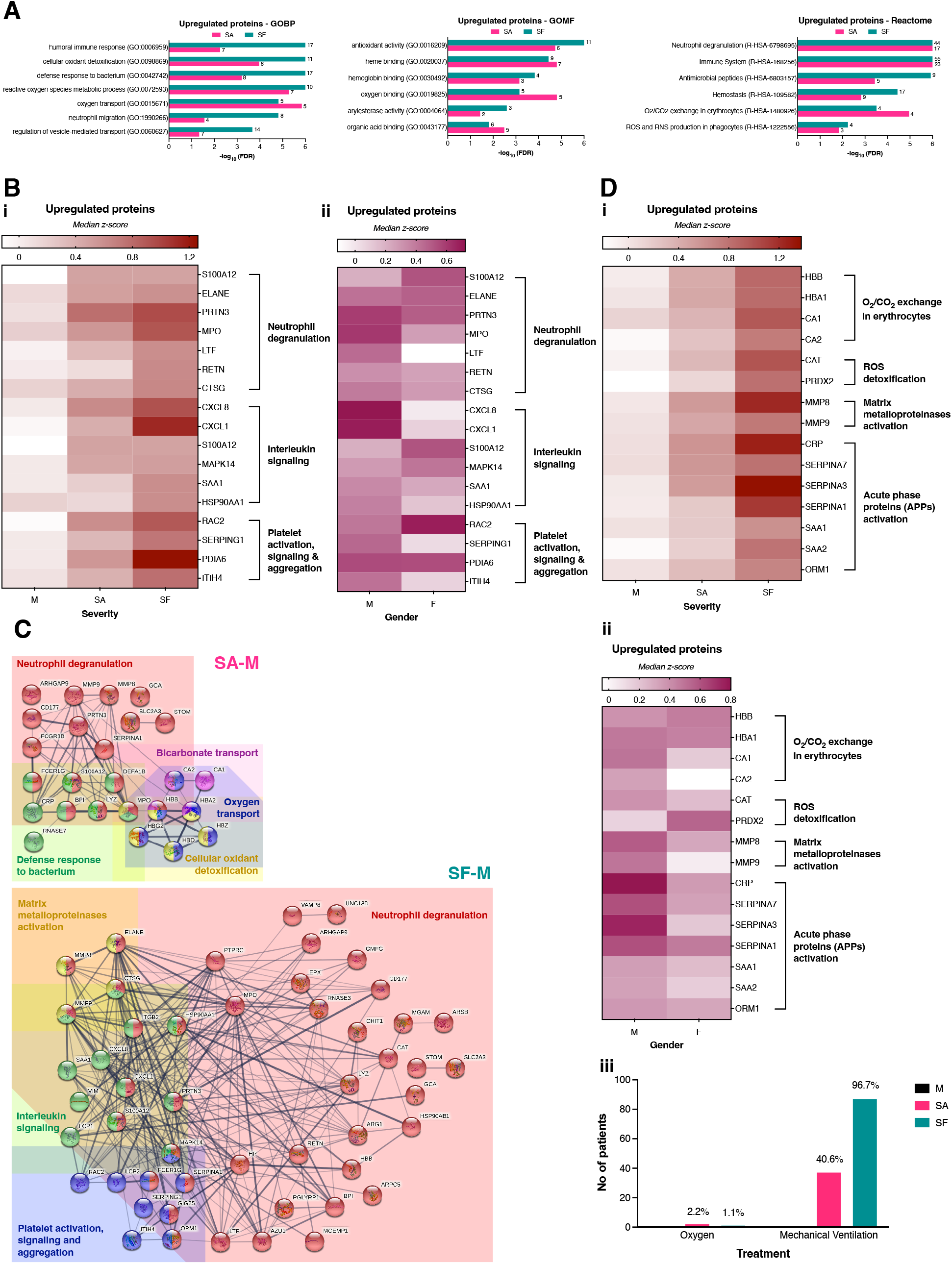
SARS-CoV-2-mediated hyperactivation of the host innate immune system in severe COVID-19 patients. **A)** Gene ontology (GO) enrichment analysis of significantly upregulated proteins (≥ 1.5 fold-change) in SA and SF groups. **B)** Upregulation of proteins involved in neutrophil degranulation, interleukin signaling, and platelet activation, signaling and aggregation pathways based on (**i**) disease severity and (**ii**) gender. **C)** Protein network analysis of upregulated proteins in SA-M and SF-M group respectively using STRING version 11.5. **D)** Effects of (**i**) disease severity and (**ii**) gender on significantly upregulated proteins in O_2_/CO_2_ gaseous exchange, detoxification of reactive oxygen species (ROS), activation of matrix metalloproteinase enzymes, and activated acute phase proteins (APPs). (**iii**) The number of ICU-admitted patients receiving oxygen support and/or mechanical ventilation. Data information: Protein abundances were plotted based on the median z-score, from the least abundant (white) to the most abundant protein level (red/violet).

The NRF2 pathway has been reported to sense the alteration of homeostasis during SARS-CoV-2 infection (Muri and Kopf, 2021). Here, we show that the NRF2 pathway is suppressed by the virus, as evidenced by the downregulation of a repertoire of NRF2-mediated-ARE-dependent antioxidant proteins such as GCLC, PRDX1, TXNRD1, AKR1C2, MGST3, NQO1, HMOX2, FTH1, and GSTA1 in severe patients (**Figure 2D** and **Figure S3B**). We found that TXNRD1 is differentially regulated between genders, with higher protein level observed in females (**Figure S3Bii)**. A higher level of NRF2 activation has been observed in female mice, providing a molecular explanation for the greater oxidative stress resistance against aging and oxidative stressors often seen in females (Rooney et al., 2018). We hypothesize that the downregulation of thioredoxin (TXNRD1) and glutaredoxin (GCLC, GSTA1 and MGST3) cellular redox systems are associated with COVID-19 pathogenesis. This is consistent with a study by (Olagnier et al., 2020) showing the downregulation of NRF2-target proteins such as HMOX1 and NQOI in SARS-CoV-2 infected Vero hTMPRSS2 cells. They also found the inhibition of NRF2 antioxidant pathway in lung biopsies of COVID-19 patients, and the expression of CXCL10, a central chemokine in the COVID-19 cytokine storm, is increased in PBMCs of severe patients. Interestingly, we also found the upregulation of CXCL8 and CXCL1 in the nasopharynx of severe patients (**Figure 3B**). Our data thereby suggest that patients with severe symptoms develop a dysregulated NRF2-mediated pathway (**Figure 2D** and **Figure S3B**), that at least in part could contribute to the impaired control of viral replication (**Figure 2Biii**) and the hyperinflammatory state (**Figure 3B** and **Figure 3D**). The use of NRF2 agonists, 4-octyl-itaconate (4-OI) and the clinically approved dimethyl fumarate (DMF) can induce a distinct IFN-independent antiviral response that is broadly effective in preventing the viral replication and suppression of SARS-CoV-2-induced inflammatory response in the host cells (Olagnier et al., 2020). Considering the dual effects of 4-OI and DMF as plausible broad-spectrum antiviral and anti-inflammatory agents, it would be interesting to investigate the therapeutic effect of NRF2 activators in severe COVID-19 patients (Alam and Czajkowsky, 2021; Bousquet et al., 2020).

We observed a dysregulation in corticosteroid biosynthesis and glucocorticoid signaling (**Figure 2Ei**) that can be associated with the aberrant hyperinflammatory response in severe COVID-19. Here, we report a decrease in the CYP11B2 expression level in severe patients, notably in SF group (**Figure 2Ei**). Interestingly, there are significant age and gender differences in the expression of CYP11B2 enzyme, with lower expression level observed in the older patients (≥ 56 years old) (**Figure 2Eii**) and higher level in females (**Figure 2Eiii**). Downregulation of CYP11B2 in turn, may cause an imbalance of mineralocorticoids or glucocorticoids, which may contribute to the unconstrained activity of host inflammatory response driving much of the severe COVID-19 pathology. Several case reports have documented the onset of adrenal insufficiency (low aldosterone level) after COVID-19 infection, suggesting a potential association between the development of clinically significant autoimmune adrenal insufficiency and COVID-19 infection (Heidarpour et al., 2020; Kumar et al., 2020; Sanchez et al., 2021). COVID-19 patients might be at risk of developing central hypocortisolism. The evidence of COVID-mediated central hypocortisolism direct towards critical illness-related corticosteroid insufficiency (CIRCI) (also known as relative adrenal insufficiency) (Pal, 2020). Commonly occurs in septic shock and ARDS, CIRCI describes glucocorticoid levels that are insufficiently low for the severity of the critical illness, resulting in a magnified systemic inflammatory response. It is also associated with a prolonged need for intensive care, and increased morbidity and mortality. Proposed mechanisms leading to CIRCI include the dysregulation of stress-responsive hypothalamic-pituitary-adrenal (HPA) axis, suppressed cortisol synthesis and metabolism, and glucocorticoid resistance (Annane et al., 2017a; Annane et al., 2017b).

We observed for the first time that the non-survivors may have developed impaired glucocorticoid receptor (GR) function, as evidenced by the increased expression levels of the chaperone protein 90 kDa heat shock protein (HSP90AB1) and the co-chaperone FK506-binding protein (FKBP52) (**Figure 2Ei**), and upregulation of SERPINA 6 (**Figure 2Ei**). This implicates a suppression in glucocorticoid availability. FKBP52 and SERPINA6 expression levels are differentially regulated by gender, with lower expression observed in females (**Figure 2Eiii**). Our data suggest that the virus can disrupt the HPA axis response in SF by suppressing the corticosteroid biosynthesis, glucocorticoid release and availability, and promotes glucocorticoid resistance through impairment of GR function (**Figure 2E**), all of which mimic the mechanisms underlying CIRCI. Consistent with our data, the randomized evaluation of COVID-19 therapy (RECOVERY) clinical trial revealed that treatment with dexamethasone, a classic synthetic glucocorticoid, enhanced the survival of severe patients (Group et al., 2021; Group et al., 2020). Patients whose disease had not progressed to a stage necessitating respiratory support had no improvement with dexamethasone, as the immunosuppressive effects of glucocorticoids at this stage of disease might hamper the antiviral responses. It is in the later stage of COVID-19 pathogenesis that the immunomodulatory effects of glucocorticoids are beneficial against the hyperinflammatory phase by disrupting the uncontrolled inflammatory feedforward loop in some patients (Cain and Cidlowski, 2020).

Severe COVID-19 pathophysiology may be driven by a dysregulation in the RAAS system (Gheblawi et al., 2020; Rysz et al., 2021). SARS-CoV-2-mediated ACE2 internalization, may cause a downregulation of ACE2 with a subsequent increase in Ang 2 level (Kuba et al., 2005), resulting in an imbalance between Ang 2 and Ang 1-7 peptide levels that can potentially aggravate the inflammatory response, typically seen in critical COVID-19. The synthesis of Ang 2, the main effector peptide of classic RAAS, begins with the cleavage of AGT into Ang 1 by renin and is subsequently converted into Ang 2 by ACE1 (**Figure 2E**). We observed an elevated protein level of AGT, notably in SF patients (**Figure 2Ei**). This increase may elevate the Ang 2 level, which may plausibly relate to severe COVID-19. By interacting with G protein-coupled receptor AT1R, Ang 2 can cause constriction of arterioles, increase in blood pressure and pulse rate, cardiac hypertrophy, and oxidative stress that promote endothelial dysfunction, vessel inflammation, thrombosis, cardiac remodeling, and insulin resistance (Cooper et al., 2021; Patel et al., 2017). While (Wu et al., 2020) reported an increased level of Ang 2 that is closely associated with lung injury in COVID-19 with no change in the renin level as compared to controls, other studies reported lower circulating plasma levels of Ang 1-7, but with normal Ang 2 level, that were associated with a more severe outcome in COVID-19 (Henry et al., 2021; Henry et al., 2020). As we observed an upregulated AGT expression (**Figure 2Ei**) and a high viral load (**Figure 2Biii**) in SF patients, we implicate that the RAAS imbalance may potentially contribute to the underlying pathophysiology of COVID-19 fatality. The use of ACE/ATR inhibitors in clinical management of COVID-19 was initially debatable (Fang et al., 2020; Jarcho et al., 2020; Mancia et al., 2020), with the hypothesis that these antagonists might increase ACE2 expression and thus, increase the SARS-Cov-2 infection risk and severity (Vaduganathan et al., 2020; Zheng et al., 2020). However, observational data now support the continued use of ACE/ATR blockers in COVID-19 patients (Flacco et al., 2020; Jung et al., 2020; Mascolo et al., 2021). Some retrospective studies have observed lower all-cause mortality in patients treated with these inhibitors (Kuster et al., 2020; Zhang et al., 2020b). A RAAS imbalance swine model mimicking several severe COVID-19 clinical presentations was developed by infusing Ang 2 or ACE2 inhibitors. These symptoms were successfully rescued using the ATR blocker and heparin (Rysz et al., 2021)). We therefore propose that SARS-CoV-2 virus manipulates the host immune system by disrupting the delicate balance between RA signaling, redox state, glucocorticoid, RAAS, and the immune system to accommodate its propagation and transmission in the host cells, rendering the COVID-19 disease progression from initial infection to severe state and eventually, fatality.

### Elevated neutrophil degranulation, interleukin signaling and platelet aggregation drive the hyperinflammatory response in severe COVID-19

We performed a functional enrichment analysis of significantly upregulated proteins and identified a prominent signature of neutrophil activation (**Figure 3** and **Figure S3C**). Studies have reported a high neutrophil-to-lymphocyte ratio (NLR), a clinical inflammation biomarker, in severe COVID-19 patients (Liu et al., 2020a; Webb et al., 2020). The increase in NLR is usually accompanied by higher D-dimer and CRP (Ponti et al., 2020; Ye et al., 2020), and we also observed the CRP increase in our severe patients (**Figure 3Di** and **Figure S3Gi**). Lung autopsies of deceased COVID-19 patients have revealed neutrophil infiltration in pulmonary capillaries, their extravasation into the alveolar spaces and neutrophilic mucositis, indicating inflammation in the entire lower respiratory tract (Barnes et al., 2020). Increased levels of circulating neutrophil extracellular traps (NETs) indicative of neutrophil activation have been described in COVID-19 (Golonka et al., 2020). NETs are composed of released neutrophil DNA and associated proteins, found in the lungs of critical COVID-19 patients, and have been shown to induce apoptosis in lung epithelial cells (Veras et al., 2020) that can contribute to immunothrombosis (Middleton et al., 2020). NETs consist of chromatin fibers associated with enzymes such as neutrophil elastase (ELANE), cathepsin G (CTSG), myeloperoxidase (MPO), and myeloblastin (PRTN3) (Delgado-Rizo et al., 2017; Kaplan and Radic, 2012; Schulte-Schrepping et al., 2020). Here, we show the significant upregulation of ELANE, CTSG, MPO, and PRTN3 in severe patients, with both CTSG and MPO expressed at higher levels in SF (**Figure 3Bi** and **Figure S3Ci**). The increased expression of ELANE, MPO and PRTN3 (**Figure 3Bi** and **Figure S3Ci**), markers of pre-matured neutrophils, suggests that severe COVID-19 is associated with the appearance of immature pre- and pro-neutrophils. The gene upregulation of *ELANE*, *MPO* and *PRTN3* has been previously shown in the pro-neutrophil cluster of PBMCs (Schulte-Schrepping et al., 2020). The increased infiltration of immature and dysfunctional neutrophils can contribute to the immune response imbalance in the lungs of severe patients. Upregulation of ELANE and MPO (**Figure 3Bi** and **Figure S3Ci**) also indicates enhanced degranulation of primary granules in the neutrophils. Similarly, high serum levels of ELANE and MPO, and increased CD63^+^ surface expression in primary granules have been found in severe COVID-19 patients (Parackova et al., 2020).

We observed significant upregulation of resistin (RETN) and lactotransferrin (LTF) (**Figure 3Bi** and **Figure S3Ci**), the neutrophil-derived effectors in SF patients. We found a lower expression of LTF in the females (**Figure S3Cii**). A single-cell atlas of the peripheral immune response has identified a developing neutrophil population expressing *LTF* in severe COVID-19 patients. These cells appear closely related to plasma blasts and are significantly enriched only in ARDS patients requiring mechanical ventilation (Wilk et al., 2020). Since RETN regulates the production of multiple cytokines (IL-6, IL-8 and TNF-*α*) (Zhang et al., 2010), the increased RETN expression promotes the release of inflammatory cytokines in SF patients, which can further exacerbate the inflammatory circuit and cause significant collateral damage to the lungs, vasculature and other organs in the non-survivors. We also found the increased expression of S100A12, a calcium-, zinc- and copper-binding protein involved in the regulation of inflammatory and immune responses in severe COVID-19 (**Figure 3Bi** and **Figure S3Ci**). Since S100A12 acts as a chemoattractant for monocytes and neutrophils, the increased expression in severe patients can stimulate neutrophil activation and degranulation and subsequently, promote cytokine production and release that further stimulates the leukocytes recruitment to the sites of inflammation. Classical monocytes and myeloid cells from severe COVID-19 patients expressed a high level of *S100A12*, the gene encoding EN-RAGE, but not the typical inflammatory mediators IL-6 and TNF-*α*. This suggests that SARS-CoV-2 infection induces a distinct inflammatory profile, as characterized by cytokines released from lung tissues but suppression of the innate immune system in the periphery (Arunachalam et al., 2020).

Indeed, we also observed hallmarks of enhanced interleukin production in severe patients (**Figure 3Bi** and **Figure S3Di**), complementing the increase in neutrophil activation and degranulation (**Figure 3**) in severe COVID-19. We found that the expression of CXCL1 and CXCL8 chemokines are associated with disease severity (**Figure 3Bi** and **Figure S3Di**) and gender (**Figure 3Bii** and **Figure S3Dii**). CXCL8 (IL-8) is a potent proinflammatory cytokine involved in the recruitment and activation of neutrophils during inflammation (Baggiolini et al., 1989). A distinct immunologic profile has been observed in the lungs of severe COVID-19 patients with ARDS, displaying a depleted and exhausted CD4 and CD8 T-cell population, but dominated by immature neutrophils, all of which reside within a milieu of proinflammatory chemokines IL-1RA, IL-6, IP-10, MCP-1 and notably, the IL-8 which is detected at very high concentrations in the lungs (Ronit et al., 2021). This suggests that IL-8 exhibits a notable compartmentalized response by recruiting neutrophils to the lungs during acute pulmonary inflammation. Hyperinflammation in the lungs of severe patients is fueled by excessive chemokines production triggering cytokine storm, as evidenced by the enriched CXCL1 and CXCL8 levels and immune cell population of neutrophils, lymphocytes and eosinophils in the bronchoalveolar lavage fluid (BALF) (Zaid et al., 2021). Upregulation of CXCL8 and CXCL1 (**Figure 3Bi** and **Figureure EV3Di**) in the nasopharynx can stimulate neutrophil activation and degranulation (**Figure 3Bi** and **Figure S3Ci**), which further promotes the release of proinflammatory cytokines and ROS (**Figure 3** and **Figure S3**) triggering cytokine storm, thereby increasing NET formation that contributes to thrombosis and viral sepsis in severe COVID-19.

Another pathophysiological mechanism linked to neutrophils in severe COVID-19 is the increase in platelet-neutrophil aggregates that promotes NET formation (Leppkes et al., 2020; Manne et al., 2020; Middleton et al., 2020). Evidences showing an association between neutrophils and activated platelets that guide their migration to damaged tissues and induce NET formation by eliciting immunothrombosis in blood vessels (Gaertner and Massberg, 2016; Hidalgo et al., 2009; Sreeramkumar et al., 2014). We show the increased expression of RAC2, SERPING1, PDIA6, and ITIH4 involved in platelet activation, signaling and aggregation (**Figure 3** and **Figure S3Ei**). The increased expression of SERPING1 is associated with disease severity (**Figure 3Bi** and **Figure S3Ei**) and gender (**Figure 3Bii** and **Figure S3Eii**). A case study has reported the increased plasma level of SERPING1 in mild and severe COVID-19 patients, implicating the role of SERPING1, a C1 esterase inhibitor, as a central regulator of contact and complement systems, potentially linking COVID-19 to complement hyperactivation, fibrin clot formation and immune depression (Hausburg et al., 2021). Elevated expression of SERPING1 could act as a negative-feedback response, an attempt by the host to inhibit contact and complement hyperactivation and hypercoagulability associated with COVID-19. We found that ITIH4, a protein belonging to the family of inter-*α*-trypsin inhibitors is significantly upregulated only in the non-survivors (**Figure S3Ei**) and males (**Figure S3Eii**). The increased ITIH4 protein expression has been found in the sera of deceased patients with severe COVID-19 (Vollmy et al., 2021) and is associated with disease severity (Messner et al., 2020). A higher level of platelet activation at rest and its interaction with neutrophils, monocytes and T cells has been observed in COVID-19 patients. The observed platelet hyperactivation, as evidenced by increased aggregation and thromboxane generation, spread on fibrinogen and collagen via upregulation of the MAPK pathway (Manne et al., 2020). Consistent with this, we show a marked increase in the expression of MAPK14 protein in severe patients (**Figure S3Di**). Another study showing severe COVID-19 patients-derived platelets exhibit increased platelet and platelet-monocyte aggregation. These platelets induce monocyte-derived tissue factor (TF) expression that can be associated with disease severity and mortality (Hottz et al., 2020). Hence, understanding the role of platelet in COVID-19-associated thrombosis has clear clinical implications in considering antiplatelet therapy, but with careful consideration taking into account the occurrence of moderate bleeding and thrombocytopenia that have already been seen in COVID-19 patients.

### Increased levels of matrix metalloproteinases (MMPs), acute phase proteins (APPs) and oxidative stress can be associated with COVID-19 mortality

We report an upregulation of MMP8 and MMP9 expression in severe patients (**Figure 3Di** and **Figure S3Fi**). We found that the expression levels of MMP8 and MMP9 proteins are lower in the female nasopharynx (**Figure 3Dii** and **Figure S3Fii**). While present in low abundance in healthy lungs, high level of MMP9 has been implicated in a wide variety of pulmonary pathologies including acute lung injury, ARDS and chronic lung diseases (Atkinson, 2003). In an acute lung injury, MMP8 and MMP9 are produced by the neutrophils, whereas the alveolar macrophages secrete MMP9, promoting inflammation and degradation of the alveolar capillary barrier. As MMPs can also act on inflammatory mediators, their release can further stimulate the migration of inflammatory cells, damaging the lung tissue (Davey et al., 2011). MMP9 was shown to be a key player contributing to the extracellular matrix remodeling and enhanced fibrosis, typically seen in COVID-19 related pneumonia. The increased plasma MMP9 strongly correlates with neutrophil counts and respiratory failure, suggesting that MMP9 can be an early indicator of respiratory failure in COVID-19 patients (Ueland et al., 2020). Moreover, MMP8, a marker of the developing neutrophil population was highly correlated with the absolute neutrophil count in severe COVID-19 patients, and its increased level is significantly associated with mortality (Meizlish et al., 2021).

As expected, we found the upregulation of APPs (CRP, SERPINA7, SERPINA3, SERPINA1, SAA1, SAA2, and ORM1) in severe patients, which are expressed at much higher levels in SF group (**Figure 3Di** and **Figure S3Gi**). These APPs are commonly associated inflammatory markers for COVID-19, which can also be reliable indicators of severe infections. Among the upregulated APPs, we found that females generally have lower expression levels of these proteins (**Figure 3Dii**), with significantly lower protein levels of CRP and SERPINA3 (**Figure S3Gii**) as compared to their male counterparts. The elevated APPs levels could potentially predispose males to hyperinflammation and poor adaptive immune responses to SARS-CoV-2 infection. SAA1, SAA2, SAA4, CRP, SERPINA3, and SAP were previously found to be significantly upregulated in the sera of severe COVID-19 patients (Gao et al., 2020; Li et al., 2020; Qin et al., 2020; Shen et al., 2020). The upregulation of CRP, SAA1, SERPINA1, SERPINA3, and SERPING1 proteins was also observed in the lung tissues and sera of patients with severe COVID-19, which could act as potential biomarkers for the degree of COVID-19-related lung damage (Wang et al., 2021).

SAA1 and ORM1 expression was reported in the lung tissues of COVID-19 patients using *in situ* immunohistofluorescence (Leng et al., 2021). The ORM1 expression is mainly controlled by a combination of the major regulatory mediators including glucocorticoids, IL-1, TNF-*α*, and IL-6, in which the increase can reach between 5 – 50 folds upon inflammation in the host cells (Baumann and Gauldie, 1990; Luo et al., 2015). A marked increase in the expression of ORM1 in SF (**Figure 3Di** and **Figure S3Gi**) can be associated with deficiency in glucocorticoid levels (**Figure 2Ei**) and upregulated interleukin signaling (**Figure 3Bi**, **Figure S3Di** and **EV4B**), implicating the involvement of these two signaling pathways in the regulation of ORM1 expression. We performed an enrichment analysis of all DE proteins for SA and SF using DisGeNET (Pinero et al., 2017) and found that the phenomena of immune-mediated thrombocytopenia and inflammation are highly enriched in SF (**Figure S4B-C**), which can be associated with ORM1 upregulation in SF (**Figure 3Di** and **Figure S3Gi**). Furthermore, we show the low level of plasminogen (PLG) expression is significantly associated with mortality (**Figure S4Ci**), inflammatory markers (**Figure 3Di** and **Figure S3Gi**), gender (**Figure S4Cii**), and age (**Figure S4Ciii**) in COVID-19 patients. The decreased PLG level in SF (**Figure S4Ci**) is consistent with a recent report demonstrating low level of PLG increases risk for mortality in COVID-19 patients, and is significantly associated with inflammatory markers including CRP, PCT, IL-6, markers of coagulation and organ dysfunctions (Della-Morte et al., 2021).

Elevated neutrophil activation (**Figure 3 and Figure S3Ci**) can trigger oxidative stress in severe patients. Interestingly, children whose neutrophils that are less reactive and adherent, with no alteration of redox balance are less prone to developing severe COVID-19 (Laforge et al., 2020). Here, we demonstrate the upregulation of catalase (CAT) and peroxiredoxin 2 (PRDX2) (**Figure 3Di** and **Figure S3H**) in severe patients. Since CAT and PRDX2 are involved in ROS detoxification, the increased oxidative stress can be associated with SARS-CoV-2 infection. It has been shown that both SOD1 and PRDX2 were notably enriched in the plasma of severe COVID-19 patients when compared to healthy controls (Xu et al., 2021). As CAT and PRDX2 are expressed at much higher levels in SF (**Figure S3H**), this suggests that oxidative stress might contribute to COVID-19 severity and mortality. The increased CAT and PRDX2 expression could not restore the cellular redox homeostasis in severe patients, suggesting that SARS-CoV-2 infection disrupts the overall antioxidant system. The RAAS system is also a key mediator of ROS production in SARS-CoV-2 infection (Alam and Czajkowsky, 2021). Ang 2 is a potent stimulator of NADPH oxidase, promoting the production of superoxide anions (O_2_^•-^) (Griendling et al., 1994; Nguyen Dinh Cat et al., 2013). The increased AGT expression in SF (**Figure 2E**) may likely reflect increased Ang 2 due to SARS-CoV-2 mediated inactivation of ACE2 (Tay et al., 2020) and subsequent predominant conversion of AGT to Ang 2, instead of Ang 1-7. Ang 2 also correlates with SARS-CoV-2 viral load (Liu et al., 2020b) and has tissue-damaging effects (Zhang et al., 2020a). The plausibility of Ang 2-induced NADPH oxidase is consistent with a report demonstrating the increased level of NADPH oxidase-induced oxidative stress in COVID-19 patients is associated with disease severity and thrombotic events (Violi et al., 2020). Earlier on, we showed that NRF2 mediated-antioxidant proteins are suppressed in severe patients (**Figure 2D** and **Figure S3B**). This suggests that respiratory viral infection is associated with decreased antioxidant defense, thus promoting oxidative damage in affected cells and tissues. With excessive ROS production, deregulated neutrophils can promote systemic inflammatory response, and improper activation of neutrophils might explain the diffuse microvascular thrombosis and capillary leak syndrome in severe COVID-19 patients (Klok et al., 2020).

Moreover, the deleterious action of ROS on the function of both pulmonary cells and red blood cells (RBC) can be viewed as a major contributor to the hypoxic respiratory failure observed in most severe COVID-19 patients (Teuwen et al., 2020). Excessive ROS accumulation can affect the RBC membrane and heme group functionality via oxidation of polyunsaturated fatty acids in the RBC membrane. This affects the diffusion of oxygen and carbon dioxide, and the deformability capability of RBC in the capillary vessels, thereby favoring thrombosis. ROS excess may also cause Fe^2+^/Fe^3+^ imbalance and disturb the iron homeostasis as iron must be maintained in the Fe^2+^ state to bind to oxygen (Laforge et al., 2020). Here, we found significant upregulation of hemoglobin subunit beta (HBB) and hemoglobin subunit alpha (HBA1) in severe patients (**Figure 3Di** and **Figure S3I**), which are markedly elevated in SF. This increase may promote efficient oxygen transport from the lung to various peripheral tissues, supplied by high oxygen level in patients receiving oxygenation and/or mechanical ventilation (**Figure 3Diii**). Nearly all (∼ 96.7 %) SF patients received mechanical ventilation, whereas 40.6 % of SA patients were mechanically ventilated (**Figure 3Diii**), which could explain the increase in hemoglobin expression. We also observed the increased expression of carbonic anhydrases 1 and 2 (CA1 and CA2) in severe patients (**Figure 3Di** and **Figure S3I**), which is higher in SF. As CA plays a crucial role in regulating many physiological processes by catalyzing the reversible hydration of CO_2_ to bicarbonate and H^+^ ions, dysregulation in the CA activity may impact the pH and CO_2_ homeostasis, thus inferring a putative role of CA in COVID-19 (Deniz et al., 2021). Elevated CA1 and CA2 expression could potentially be a body response to excessive CO_2_ resulted from SARS-CoV-2-induced damaged lung cells, notably in severe patients associated with pneumonia or ARDS. CA can also modify the affinity of hemoglobin for oxygen (Gai et al., 2003). The increase in CA activity could impact the pH in erythrocytes that further disrupts the ability of hemoglobin to bind to oxygen. Hence, the use of CA inhibitors in treating severe COVID-19-related respiratory abnormalities could help to restore the balance between CO_2_ and HCO_3_^-^ levels. For example, acetazolamide has been proposed to improve the respiratory conditions in severe COVID-19 (Solaimanzadeh, 2020) through its compensatory hyperventilation with increased blood O_2_ levels and decreased blood CO_2_ levels (Leaf and Goldfarb, 2007).

Overall, we demonstrate the proteome signatures capturing the host response to COVID-19 infection in the nasopharynx, highlighting that severe COVID-19 is a double-defect disease, requiring first a defective early control of the virus mainly through the 1) IFN-I depletion via aberrant STAT1 signaling, retinol metabolism, NRF2 antioxidant system, and dysregulated glucocorticoid and RAAS signaling, and 2) an impaired ability to control proinflammatory responses by hyperactivating the host innate immunological pathways through neutrophil activation and degranulation, interleukin production and platelet aggregation (**Figure 4**). Only when both of these checkpoints fail will the pathogenesis of severe COVID-19 develop, or worse, when the perpetuation of self-sustained inflammatory circuits will lead to fatality. With the feasibility of NP swab as routinely used clinical sample for detecting COVID-19, we believe that the identified signatures can act as clinical classifiers, paving way for the development of routine assays to support clinical decision making. Our data also provide resources for mining potential COVID-19 therapeutic targets, which warrant further validation on larger-scale and longitudinal studies, in parallel with a compendium of standard accredited clinical tests to capture the diverse disease trajectories.

**Figure 4.**
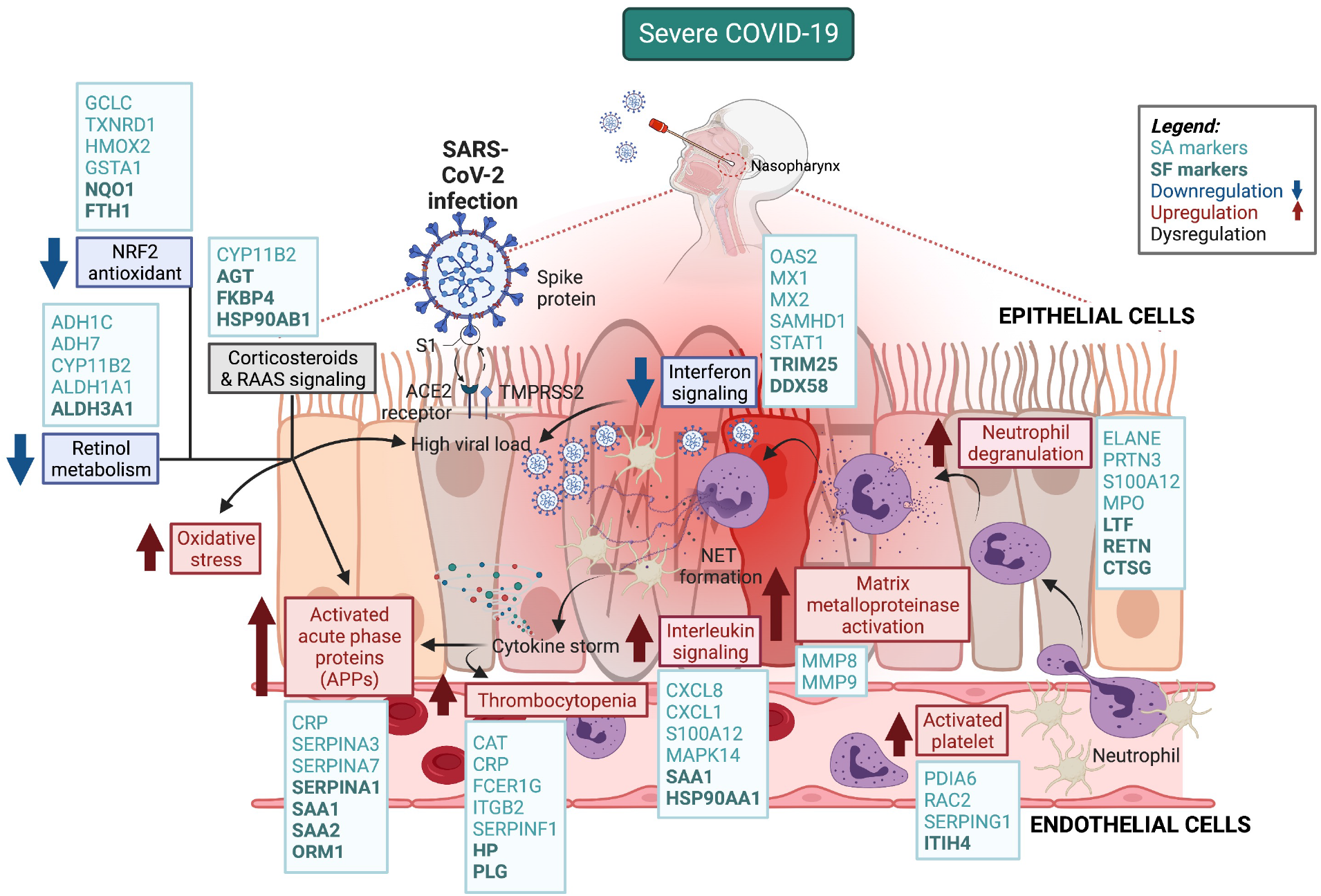
Protein markers and their associated pathways that are unique to the pathophysiology of COVID-19 disease severity and fatality. SARS-CoV-2 virus hijacks the host innate immune system to accommodate its replication by impairing the STAT1-mediated IFN-I response, and depletes the host retinol metabolism and NRF2 antioxidant system that are associated with disease severity. The infection also dysregulates the glucocorticoid signaling and the renin-angiotensin-aldosterone system (RAAS) which can contribute to the pathophysiology of COVID-19 fatality. This virus also hyperactivates the host innate immune system through neutrophil activation and degranulation, interleukin production and platelet aggregation that leads to cytokine storm, coagulopathy and thrombocytopenia in severe patients. These proteome signatures capturing the host response to COVID-19 infection in the nasopharynx highlight that severe COVID-19 is a double-defect disease, requiring a defective early control of the virus and an impaired ability to control proinflammatory responses.

## Limitations of the study

We could not match the exact number of patients for each age and gender category to severity. This is due to a strict lockdown imposed by the authorities in Saudi Arabia during the pandemic, and the participating hospitals received more male patients, thereby restricting our access to female samples. Furthermore, we were unable to enroll healthy subjects as control at that point of our study, hence, we did not include and match negative healthy controls to our COVID-19 positive cases.

## Acknowledgements

We sincerely thank all the hospital staff members, including the team from the General Directorate of Health Affairs in the Makkah region for their tremendous help in collecting the nasopharyngeal swab samples and patients’ metadata. We also thank the KAUST Health Safety and Environment (HSE) and Security staff for their support in logistics relating to sample transport during the COVID-19 lockdown in KSA in 2020.

## Author Contributions

Conceptualization, A.P.^1^, A.O., and L.E.E.; Methodology, A.O. and L.E.E; Investigation, A.O, L.E.E, A.P.^2^, A.G., S.M., R.P.S., A.K.S, and F.B.R.; Formal Analysis, A.P.^2^ and A.O.; Resources, F.A., A.S., K.A., A.K., A.H., N.A., and S.H.; Visualization, A.O. and A.P.^2^; Writing – Original Draft, A.O., L.E.E., A.P.^2^, and A.G.; Writing – Review & Editing, A.P.^1^, P.J.M., A.O., and L.E.E.; Supervision, A.P.^1^ and P.J.M.; Funding Acquisition, A.P.^1^

## References

Alam, M.S., and Czajkowsky, D.M. (2021). SARS-CoV-2 infection and oxidative stress: Pathophysiological insight into thrombosis and therapeutic opportunities. Cytokine Growth Factor Rev.

Alwan, N.A., Burgess, R.A., Ashworth, S., Beale, R., Bhadelia, N., Bogaert, D., Dowd, J., Eckerle, I., Goldman, L.R., Greenhalgh, T., et al. (2020). Scientific consensus on the COVID-19 pandemic: we need to act now. Lancet 396, e71–e72.

Andolfo, I., Russo, R., Lasorsa, V.A., Cantalupo, S., Rosato, B.E., BonFigurelio, F., Frisso, G., Abete, P., Cassese, G.M., Servillo, G., et al. (2021). Common variants at 21q22.3 locus influence MX1 and TMPRSS2 gene expression and susceptibility to severe COVID-19. iScience 24, 102322.

Annane, D., Pastores, S.M., Arlt, W., Balk, R.A., Beishuizen, A., Briegel, J., Carcillo, J., Christ-Crain, M., Cooper, M.S., Marik, P.E., et al. (2017a). Critical illness-related corticosteroid insufficiency (CIRCI): a narrative review from a Multispecialty Task Force of the Society of Critical Care Medicine (SCCM) and the European Society of Intensive Care Medicine (ESICM). Intensive Care Med 43, 1781–1792.

Annane, D., Pastores, S.M., Rochwerg, B., Arlt, W., Balk, R.A., Beishuizen, A., Briegel, J., Carcillo, J., Christ-Crain, M., Cooper, M.S., et al. (2017b). Guidelines for the Diagnosis and Management of Critical Illness-Related Corticosteroid Insufficiency (CIRCI) in Critically Ill Patients (Part I): Society of Critical Care Medicine (SCCM) and European Society of Intensive Care Medicine (ESICM) 2017. Crit Care Med 45, 2078–2088.

Arunachalam, P.S., Wimmers, F., Mok, C.K.P., Perera, R., Scott, M., Hagan, T., Sigal, N., Feng, Y., Bristow, L., Tak-Yin Tsang, O., et al. (2020). Systems biological assessment of immunity to mild versus severe COVID-19 infection in humans. Science 369, 1210–1220.

Asano, T., Boisson, B., Onodi, F., Matuozzo, D., Moncada-Velez, M., Maglorius Renkilaraj, M.R.L., Zhang, P., Meertens, L., Bolze, A., Materna, M., et al. (2021). X-linked recessive TLR7 deficiency in ∼1% of men under 60 years old with life-threatening COVID-19. Sci Immunol 6.

Atkinson, S. (2003). The HSJ interview: Dr Sue Atkinson. Double impact. Interview by Helen Mooney. Health Serv J 113, 22–23.

Baggiolini, M., Walz, A., and Kunkel, S.L. (1989). Neutrophil-activating peptide-1/interleukin 8, a novel cytokine that activates neutrophils. J Clin Invest 84, 1045–1049.

Barnes, B.J., Adrover, J.M., Baxter-Stoltzfus, A., Borczuk, A., Cools-Lartigue, J., Crawford, J.M., Dassler-Plenker, J., Guerci, P., Huynh, C., Knight, J.S., et al. (2020). Targeting potential drivers of COVID-19: Neutrophil extracellular traps. J Exp Med 217.

Bastard, P., Rosen, L.B., Zhang, Q., Michailidis, E., Hoffmann, H.H., Zhang, Y., Dorgham, K., Philippot, Q., Rosain, J., Beziat, V., et al. (2020). Autoantibodies against type I IFNs in patients with life-threatening COVID-19. Science 370.

Baumann, H., and Gauldie, J. (1990). Regulation of hepatic acute phase plasma protein genes by hepatocyte stimulating factors and other mediators of inflammation. Mol Biol Med 7, 147–159.

Bhargava, A., Lahaye, X., and Manel, N. (2018). Let me in: Control of HIV nuclear entry at the nuclear envelope. Cytokine Growth Factor Rev 40, 59–67.

Bizzotto, J., Sanchis, P., Abbate, M., Lage-Vickers, S., Lavignolle, R., Toro, A., Olszevicki, S., Sabater, A., Cascardo, F., Vazquez, E., et al. (2020). SARS-CoV-2 Infection Boosts MX1 Antiviral Effector in COVID-19 Patients. iScience 23, 101585.

Blanco-Melo, D., Nilsson-Payant, B.E., Liu, W.C., Uhl, S., Hoagland, D., Moller, R., Jordan, T.X., Oishi, K., Panis, M., Sachs, D., et al. (2020). Imbalanced Host Response to SARS-CoV-2 Drives Development of COVID-19. Cell 181, 1036–1045 e1039.

Blumenthal, D., Fowler, E.J., Abrams, M., and Collins, S.R. (2020). Covid-19 - Implications for the Health Care System. N Engl J Med 383, 1483–1488.

Bousquet, J., Cristol, J.P., Czarlewski, W., Anto, J.M., Martineau, A., Haahtela, T., Fonseca, S.C., Iaccarino, G., Blain, H., Fiocchi, A., et al. (2020). Nrf2-interacting nutrients and COVID-19: time for research to develop adaptation strategies. Clin Transl Allergy 10, 58.

Bunders, M.J., and Altfeld, M. (2020). Implications of Sex Differences in Immunity for SARS-CoV-2 Pathogenesis and Design of Therapeutic Interventions. Immunity 53, 487–495.

Cain, D.W., and Cidlowski, J.A. (2020). After 62 years of regulating immunity, dexamethasone meets COVID-19. Nat Rev Immunol 20, 587–588.

Casadevall, A., and Pirofski, L.A. (2020). In fatal COVID-19, the immune response can control the virus but kill the patient. Proc Natl Acad Sci U S A 117, 30009–30011.

Chattha, K.S., Kandasamy, S., Vlasova, A.N., and Saif, L.J. (2013). Vitamin A deficiency impairs adaptive B and T cell responses to a prototype monovalent attenuated human rotavirus vaccine and virulent human rotavirus challenge in a gnotobiotic piglet model. PLoS One 8, e82966.

Chen, Z., Lin, F., Gao, Y., Li, Z., Zhang, J., Xing, Y., Deng, Z., Yao, Z., Tsun, A., and Li, B. (2011). FOXP3 and RORgammat: transcriptional regulation of Treg and Th17. Int Immunopharmacol 11, 536–542.

Chow, K.T., Gale, M., Jr., and Loo, Y.M. (2018). RIG-I and Other RNA Sensors in Antiviral Immunity. Annu Rev Immunol 36, 667–694.

Combes, A.J., Courau, T., Kuhn, N.F., Hu, K.H., Ray, A., Chen, W.S., Chew, N.W., Cleary, S.J., Kushnoor, D., Reeder, G.C., et al. (2021). Global absence and targeting of protective immune states in severe COVID-19. Nature 591, 124–130.

Cooper, S.L., Boyle, E., Jefferson, S.R., Heslop, C.R.A., Mohan, P., Mohanraj, G.G.J., Sidow, H.A., Tan, R.C.P., Hill, S.J., and Woolard, J. (2021). Role of the Renin-Angiotensin-Aldosterone and Kinin-Kallikrein Systems in the Cardiovascular Complications of COVID-19 and Long COVID. Int J Mol Sci 22.

Dalskov, L., Mohlenberg, M., Thyrsted, J., Blay-Cadanet, J., Poulsen, E.T., Folkersen, B.H., Skaarup, S.H., Olagnier, D., Reinert, L., Enghild, J.J., et al. (2020). SARS-CoV-2 evades immune detection in alveolar macrophages. EMBO Rep 21, e51252.

Davey, A., McAuley, D.F., and O’Kane, C.M. (2011). Matrix metalloproteinases in acute lung injury: mediators of injury and drivers of repair. Eur Respir J 38, 959–970.

Del Valle, D.M., Kim-Schulze, S., Huang, H.H., Beckmann, N.D., Nirenberg, S., Wang, B., Lavin, Y., Swartz, T.H., Madduri, D., Stock, A., et al. (2020). An inflammatory cytokine signature predicts COVID-19 severity and survival. Nat Med 26, 1636–1643.

Delgado-Rizo, V., Martinez-Guzman, M.A., Iniguez-Gutierrez, L., Garcia-Orozco, A., Alvarado-Navarro, A., and Fafutis-Morris, M. (2017). Neutrophil Extracellular Traps and Its Implications in Inflammation: An Overview. Front Immunol 8, 81.

Della-Morte, D., Pacifici, F., Ricordi, C., Massoud, R., Rovella, V., Proietti, S., Iozzo, M., Lauro, D., Bernardini, S., Bonassi, S., et al. (2021). Low level of plasminogen increases risk for mortality in COVID-19 patients. Cell Death Dis 12, 773.

Deniz, S., Uysal, T.K., Capasso, C., Supuran, C.T., and Ozensoy Guler, O. (2021). Is carbonic anhydrase inhibition useful as a complementary therapy of Covid-19 infection? J Enzyme Inhib Med Chem 36, 1230–1235.

Dorward, D.A., Russell, C.D., Um, I.H., Elshani, M., Armstrong, S.D., Penrice-Randal, R., Millar, T., Lerpiniere, C.E.B., Tagliavini, G., Hartley, C.S., et al. (2021). Tissue-Specific Immunopathology in Fatal COVID-19. Am J Respir Crit Care Med 203, 192–201.

Elias, K.M., Laurence, A., Davidson, T.S., Stephens, G., Kanno, Y., Shevach, E.M., and O’Shea, J.J. (2008). Retinoic acid inhibits Th17 polarization and enhances FoxP3 expression through a Stat-3/Stat-5 independent signaling pathway. Blood 111, 1013–1020.

Fang, L., Karakiulakis, G., and Roth, M. (2020). Are patients with hypertension and diabetes mellitus at increased risk for COVID-19 infection? Lancet Respir Med 8, e21.

Flacco, M.E., Acuti Martellucci, C., Bravi, F., Parruti, G., Cappadona, R., Mascitelli, A., Manfredini, R., Mantovani, L.G., and Manzoli, L. (2020). Treatment with ACE inhibitors or ARBs and risk of severe/lethal COVID-19: a meta-analysis. Heart 106, 1519–1524.

Gack, M.U., Shin, Y.C., Joo, C.H., Urano, T., Liang, C., Sun, L., Takeuchi, O., Akira, S., Chen, Z., Inoue, S., et al. (2007). TRIM25 RING-finger E3 ubiquitin ligase is essential for RIG-I-mediated antiviral activity. Nature 446, 916–920.

Gaertner, F., and Massberg, S. (2016). Blood coagulation in immunothrombosis-At the frontline of intravascular immunity. Semin Immunol 28, 561–569.

Gai, X., Taki, K., Kato, H., and Nagaishi, H. (2003). Regulation of hemoglobin affinity for oxygen by carbonic anhydrase. J Lab Clin Med 142, 414–420.

Galani, I.E., Rovina, N., Lampropoulou, V., Triantafyllia, V., Manioudaki, M., Pavlos, E., Koukaki, E., Fragkou, P.C., Panou, V., Rapti, V., et al. (2021). Untuned antiviral immunity in COVID-19 revealed by temporal type I/III interferon patterns and flu comparison. Nat Immunol 22, 32–40.

Gao, Y., Li, T., Han, M., Li, X., Wu, D., Xu, Y., Zhu, Y., Liu, Y., Wang, X., and Wang, L. (2020). Diagnostic utility of clinical laboratory data determinations for patients with the severe COVID-19. J Med Virol 92, 791–796.

Gheblawi, M., Wang, K., Viveiros, A., Nguyen, Q., Zhong, J.C., Turner, A.J., Raizada, M.K., Grant, M.B., and Oudit, G.Y. (2020). Angiotensin-Converting Enzyme 2: SARS-CoV-2 Receptor and Regulator of the Renin-Angiotensin System: Celebrating the 20th Anniversary of the Discovery of ACE2. Circ Res 126, 1456–1474.

Golonka, R.M., Saha, P., Yeoh, B.S., Chattopadhyay, S., Gewirtz, A.T., Joe, B., and Vijay-Kumar, M. (2020). Harnessing innate immunity to eliminate SARS-CoV-2 and ameliorate COVID-19 disease. Physiol Genomics 52, 217–221.

Gori Savellini, G., Anichini, G., Gandolfo, C., and Cusi, M.G. (2021). SARS-CoV-2 N Protein Targets TRIM25-Mediated RIG-I Activation to Suppress Innate Immunity. Viruses 13.

Grant, R.A., Morales-Nebreda, L., Markov, N.S., Swaminathan, S., Querrey, M., Guzman, E.R., Abbott, D.A., Donnelly, H.K., Donayre, A., Goldberg, I.A., et al. (2021). Circuits between infected macrophages and T cells in SARS-CoV-2 pneumonia. Nature 590, 635–641.

Griendling, K.K., Minieri, C.A., Ollerenshaw, J.D., and Alexander, R.W. (1994). Angiotensin II stimulates NADH and NADPH oxidase activity in cultured vascular smooth muscle cells. Circ Res 74, 1141–1148.

Group, R.C., Horby, P., Lim, W.S., Emberson, J.R., Mafham, M., Bell, J.L., Linsell, L., Staplin, N., Brightling, C., Ustianowski, A., et al. (2021). Dexamethasone in Hospitalized Patients with Covid-19. N Engl J Med 384, 693–704.

Group, W.H.O.R.E.A.f.C.-T.W., Sterne, J.A.C., Murthy, S., Diaz, J.V., Slutsky, A.S., Villar, J., Angus, D.C., Annane, D., Azevedo, L.C.P., Berwanger, O., et al. (2020). Association Between Administration of Systemic Corticosteroids and Mortality Among Critically Ill Patients With COVID-19: A Meta-analysis. JAMA 324, 1330–1341.

Guarda, G., Braun, M., Staehli, F., Tardivel, A., Mattmann, C., Forster, I., Farlik, M., Decker, T., Du Pasquier, R.A., Romero, P., et al. (2011). Type I interferon inhibits interleukin-1 production and inflammasome activation. Immunity 34, 213–223.

Hadjadj, J., Yatim, N., Barnabei, L., Corneau, A., Boussier, J., Smith, N., Pere, H., Charbit, B., Bondet, V., Chenevier-Gobeaux, C., et al. (2020). Impaired type I interferon activity and inflammatory responses in severe COVID-19 patients. Science 369, 718–724.

Haller, O., Staeheli, P., Schwemmle, M., and Kochs, G. (2015). Mx GTPases: dynamin-like antiviral machines of innate immunity. Trends Microbiol 23, 154–163.

Harrison, A.G., Lin, T., and Wang, P. (2020). Mechanisms of SARS-CoV-2 Transmission and Pathogenesis. Trends Immunol 41, 1100–1115.

Hausburg, M.A., Banton, K.L., Roshon, M., and Bar-Or, D. (2021). Clinically distinct COVID-19 cases share notably similar immune response progression: A follow-up analysis. Heliyon 7, e05877.

Heidarpour, M., Vakhshoori, M., Abbasi, S., Shafie, D., and Rezaei, N. (2020). Adrenal insufficiency in coronavirus disease 2019: a case report. J Med Case Rep 14, 134.

Henry, B.M., Benoit, J.L., Berger, B.A., Pulvino, C., Lavie, C.J., Lippi, G., and Benoit, S.W. (2021). Coronavirus disease 2019 is associated with low circulating plasma levels of angiotensin 1 and angiotensin 1,7. J Med Virol 93, 678–680.

Henry, B.M., Benoit, S., Lippi, G., and Benoit, J. (2020). Letter to the Editor - Circulating plasma levels of angiotensin II and aldosterone in patients with coronavirus disease 2019 (COVID-19): A preliminary report. Prog Cardiovasc Dis 63, 702–703.

Hidalgo, A., Chang, J., Jang, J.E., Peired, A.J., Chiang, E.Y., and Frenette, P.S. (2009). Heterotypic interactions enabled by polarized neutrophil microdomains mediate thromboinflammatory injury. Nat Med 15, 384–391.

Hottz, E.D., Azevedo-Quintanilha, I.G., Palhinha, L., Teixeira, L., Barreto, E.A., Pao, C.R.R., Righy, C., Franco, S., Souza, T.M.L., Kurtz, P., et al. (2020). Platelet activation and platelet-monocyte aggregate formation trigger tissue factor expression in patients with severe COVID-19. Blood 136, 1330–1341.

Huang, C., Wang, Y., Li, X., Ren, L., Zhao, J., Hu, Y., Zhang, L., Fan, G., Xu, J., Gu, X., et al. (2020). Clinical features of patients infected with 2019 novel coronavirus in Wuhan, China. Lancet 395, 497–506.

Hughes, C.S., Moggridge, S., Muller, T., Sorensen, P.H., Morin, G.B., and Krijgsveld, J. (2019). Single-pot, solid-phase-enhanced sample preparation for proteomics experiments. Nat Protoc 14, 68–85.

Itoh, K., Ishii, T., Wakabayashi, N., and Yamamoto, M. (1999). Regulatory mechanisms of cellular response to oxidative stress. Free Radic Res 31, 319–324.

Jarcho, J.A., Ingelfinger, J.R., Hamel, M.B., D’Agostino, R.B., Sr., and Harrington, D.P. (2020). Inhibitors of the Renin-Angiotensin-Aldosterone System and Covid-19. N Engl J Med 382, 2462–2464.

Jung, S.Y., Choi, J.C., You, S.H., and Kim, W.Y. (2020). Association of Renin-angiotensin-aldosterone System Inhibitors With Coronavirus Disease 2019 (COVID-19)-Related Outcomes in Korea: A Nationwide Population-based Cohort Study. Clin Infect Dis 71, 2121–2128.

Kaplan, M.J., and Radic, M. (2012). Neutrophil extracellular traps: double-edged swords of innate immunity. J Immunol 189, 2689–2695.

Kawai, T., and Akira, S. (2008). Toll-like receptor and RIG-I-like receptor signaling. Ann N Y Acad Sci 1143, 1–20.

Kawai, T., and Akira, S. (2010). The role of pattern-recognition receptors in innate immunity: update on Toll-like receptors. Nat Immunol 11, 373–384.

Kim, C.H. (2008). Regulation of FoxP3 regulatory T cells and Th17 cells by retinoids. Clin Dev Immunol 2008, 416910.

King, C., and Sprent, J. (2021). Dual Nature of Type I Interferons in SARS-CoV-2-Induced Inflammation. Trends Immunol 42, 312–322.

Klein, S.L., and Flanagan, K.L. (2016). Sex differences in immune responses. Nat Rev Immunol 16, 626–638.

Klok, F.A., Kruip, M., van der Meer, N.J.M., Arbous, M.S., Gommers, D., Kant, K.M., Kaptein, F.H.J., van Paassen, J., Stals, M.A.M., Huisman, M.V., et al. (2020). Confirmation of the high cumulative incidence of thrombotic complications in critically ill ICU patients with COVID-19: An updated analysis. Thromb Res 191, 148–150.

Kobayashi, E.H., Suzuki, T., Funayama, R., Nagashima, T., Hayashi, M., Sekine, H., Tanaka, N., Moriguchi, T., Motohashi, H., Nakayama, K., et al. (2016). Nrf2 suppresses macrophage inflammatory response by blocking proinflammatory cytokine transcription. Nat Commun 7, 11624.

Komaravelli, N., and Casola, A. (2014). Respiratory Viral Infections and Subversion of Cellular Antioxidant Defenses. J Pharmacogenomics Pharmacoproteomics 5.

Kuba, K., Imai, Y., Rao, S., Gao, H., Guo, F., Guan, B., Huan, Y., Yang, P., Zhang, Y., Deng, W., et al. (2005). A crucial role of angiotensin converting enzyme 2 (ACE2) in SARS coronavirus-induced lung injury. Nat Med 11, 875–879.

Kuleshov, M.V., Jones, M.R., Rouillard, A.D., Fernandez, N.F., Duan, Q., Wang, Z., Koplev, S., Jenkins, S.L., Jagodnik, K.M., Lachmann, A., et al. (2016). Enrichr: a comprehensive gene set enrichment analysis web server 2016 update. Nucleic Acids Res 44, W90–97.

Kumar, R., Guruparan, T., Siddiqi, S., Sheth, R., Jacyna, M., Naghibi, M., and Vrentzou, E. (2020). A case of adrenal infarction in a patient with COVID 19 infection. BJR Case Rep 6, 20200075.

Kuster, G.M., Pfister, O., Burkard, T., Zhou, Q., Twerenbold, R., Haaf, P., Widmer, A.F., and Osswald, S. (2020). SARS-CoV2: should inhibitors of the renin-angiotensin system be withdrawn in patients with COVID-19? Eur Heart J 41, 1801–1803.

Laforge, M., Elbim, C., Frere, C., Hemadi, M., Massaad, C., Nuss, P., Benoliel, J.J., and Becker, C. (2020). Author Correction: Tissue damage from neutrophil-induced oxidative stress in COVID-19. Nat Rev Immunol 20, 579.

Le Page, C., Genin, P., Baines, M.G., and Hiscott, J. (2000). Interferon activation and innate immunity. Rev Immunogenet 2, 374–386.

Leaf, D.E., and Goldfarb, D.S. (2007). Mechanisms of action of acetazolamide in the prophylaxis and treatment of acute mountain sickness. J Appl Physiol (1985) 102, 1313–1322.

Lee, J.S., and Shin, E.C. (2020). The type I interferon response in COVID-19: implications for treatment. Nat Rev Immunol 20, 585–586.

Lei, X., Dong, X., Ma, R., Wang, W., Xiao, X., Tian, Z., Wang, C., Wang, Y., Li, L., Ren, L., et al. (2020). Activation and evasion of type I interferon responses by SARS-CoV-2. Nat Commun 11, 3810.

Leng, L., Li, M., Li, W., Mou, D., Liu, G., Ma, J., Zhang, S., Li, H., Cao, R., and Zhong, W. (2021). Sera proteomic features of active and recovered COVID-19 patients: potential diagnostic and prognostic biomarkers. Signal Transduct Target Ther 6, 216.

Leppkes, M., Knopf, J., Naschberger, E., Lindemann, A., Singh, J., Herrmann, I., Sturzl, M., Staats, L., Mahajan, A., Schauer, C., et al. (2020). Vascular occlusion by neutrophil extracellular traps in COVID-19. EBioMedicine 58, 102925.

Li, H., Xiang, X., Ren, H., Xu, L., Zhao, L., Chen, X., Long, H., Wang, Q., and Wu, Q. (2020). Serum Amyloid A is a biomarker of severe Coronavirus Disease and poor prognosis. J Infect 80, 646–655.

Liu, J., Liu, Y., Xiang, P., Pu, L., Xiong, H., Li, C., Zhang, M., Tan, J., Xu, Y., Song, R., et al. (2020a). Neutrophil-to-lymphocyte ratio predicts critical illness patients with 2019 coronavirus disease in the early stage. J Transl Med 18, 206.

Liu, Y., Yang, Y., Zhang, C., Huang, F., Wang, F., Yuan, J., Wang, Z., Li, J., Li, J., Feng, C., et al. (2020b). Clinical and biochemical indexes from 2019-nCoV infected patients linked to viral loads and lung injury. Sci China Life Sci 63, 364–374.

Lu, L., Lan, Q., Li, Z., Zhou, X., Gu, J., Li, Q., Wang, J., Chen, M., Liu, Y., Shen, Y., et al. (2014). Critical role of all-trans retinoic acid in stabilizing human natural regulatory T cells under inflammatory conditions. Proc Natl Acad Sci U S A 111, E3432–3440.

Lu, X., Wang, L., Sakthivel, S.K., Whitaker, B., Murray, J., Kamili, S., Lynch, B., Malapati, L., Burke, S.A., Harcourt, J., et al. (2020). US CDC Real-Time Reverse Transcription PCR Panel for Detection of Severe Acute Respiratory Syndrome Coronavirus 2. Emerg Infect Dis 26.

Luo, Z., Lei, H., Sun, Y., Liu, X., and Su, D.F. (2015). Orosomucoid, an acute response protein with multiple modulating activities. J Physiol Biochem 71, 329–340.

Mancia, G., Rea, F., Ludergnani, M., Apolone, G., and Corrao, G. (2020). Renin-Angiotensin-Aldosterone System Blockers and the Risk of Covid-19. N Engl J Med 382, 2431–2440.

Manne, B.K., Denorme, F., Middleton, E.A., Portier, I., Rowley, J.W., Stubben, C., Petrey, A.C., Tolley, N.D., Guo, L., Cody, M., et al. (2020). Platelet gene expression and function in patients with COVID-19. Blood 136, 1317–1329.

Mascolo, A., Scavone, C., Rafaniello, C., De Angelis, A., Urbanek, K., di Mauro, G., Cappetta, D., Berrino, L., Rossi, F., and Capuano, A. (2021). The Role of Renin-Angiotensin-Aldosterone System in the Heart and Lung: Focus on COVID-19. Front Pharmacol 12, 667254.

Masood, K.I., Yameen, M., Ashraf, J., Shahid, S., Mahmood, S.F., Nasir, A., Nasir, N., Jamil, B., Ghanchi, N.K., Khanum, I., et al. (2021). Upregulated type I interferon responses in asymptomatic COVID-19 infection are associated with improved clinical outcome. Sci Rep 11, 22958.

Matsuyama, T., Kubli, S.P., Yoshinaga, S.K., Pfeffer, K., and Mak, T.W. (2020). An aberrant STAT pathway is central to COVID-19. Cell Death Differ 27, 3209–3225.

Mehta, P., McAuley, D.F., Brown, M., Sanchez, E., Tattersall, R.S., Manson, J.J., and Hlh Across Speciality Collaboration, U.K. (2020). COVID-19: consider cytokine storm syndromes and immunosuppression. Lancet 395, 1033–1034.

Meizlish, M.L., Pine, A.B., Bishai, J.D., Goshua, G., Nadelmann, E.R., Simonov, M., Chang, C.H., Zhang, H., Shallow, M., Bahel, P., et al. (2021). A neutrophil activation signature predicts critical illness and mortality in COVID-19. Blood Adv 5, 1164–1177.

Messner, C.B., Demichev, V., Wendisch, D., Michalick, L., White, M., Freiwald, A., Textoris-Taube, K., Vernardis, S.I., Egger, A.S., Kreidl, M., et al. (2020). Ultra-High-Throughput Clinical Proteomics Reveals Classifiers of COVID-19 Infection. Cell Syst 11, 11–24 e14.

Mi, H., Ebert, D., Muruganujan, A., Mills, C., Albou, L.P., Mushayamaha, T., and Thomas, P.D. (2021). PANTHER version 16: a revised family classification, tree-based classification tool, enhancer regions and extensive API. Nucleic Acids Res 49, D394–D403.

Middleton, E.A., He, X.Y., Denorme, F., Campbell, R.A., Ng, D., Salvatore, S.P., Mostyka, M., Baxter-Stoltzfus, A., Borczuk, A.C., Loda, M., et al. (2020). Neutrophil extracellular traps contribute to immunothrombosis in COVID-19 acute respiratory distress syndrome. Blood 136, 1169–1179.

Mogensen, T.H. (2009). Pathogen recognition and inflammatory signaling in innate immune defenses. Clin Microbiol Rev 22, 240–273, Table of Contents.

Mu, J., Fang, Y., Yang, Q., Shu, T., Wang, A., Huang, M., Jin, L., Deng, F., Qiu, Y., and Zhou, X. (2020). SARS-CoV-2 N protein antagonizes type I interferon signaling by suppressing phosphorylation and nuclear translocation of STAT1 and STAT2. Cell Discov 6, 65.

Mun, D.G., Vanderboom, P.M., Madugundu, A.K., Garapati, K., Chavan, S., Peterson, J.A., Saraswat, M., and Pandey, A. (2021). DIA-Based Proteome Profiling of Nasopharyngeal Swabs from COVID-19 Patients. J Proteome Res 20, 4165–4175.

Muri, J., and Kopf, M. (2021). Redox regulation of immunometabolism. Nat Rev Immunol 21, 363–381.

Naqvi, A.A.T., Fatima, K., Mohammad, T., Fatima, U., Singh, I.K., Singh, A., Atif, S.M., Hariprasad, G., Hasan, G.M., and Hassan, M.I. (2020). Insights into SARS-CoV-2 genome, structure, evolution, pathogenesis and therapies: Structural genomics approach. Biochim Biophys Acta Mol Basis Dis 1866, 165878.

Nguyen Dinh Cat, A., Montezano, A.C., Burger, D., and Touyz, R.M. (2013). Angiotensin II, NADPH oxidase, and redox signaling in the vasculature. Antioxid Redox Signal 19, 1110–1120.

Olagnier, D., Farahani, E., Thyrsted, J., Blay-Cadanet, J., Herengt, A., Idorn, M., Hait, A., Hernaez, B., Knudsen, A., Iversen, M.B., et al. (2020). SARS-CoV2-mediated suppression of NRF2-signaling reveals potent antiviral and anti-inflammatory activity of 4-octyl-itaconate and dimethyl fumarate. Nat Commun 11, 4938.

Onoguchi, K., Yoneyama, M., and Fujita, T. (2011). Retinoic acid-inducible gene-I-like receptors. J Interferon Cytokine Res 31, 27–31.

Pal, R. (2020). COVID-19, hypothalamo-pituitary-adrenal axis and clinical implications. Endocrine 68, 251–252.

Paludan, S.R., and Mogensen, T.H. (2022). Innate immunological pathways in COVID-19 pathogenesis. Sci Immunol 7, eabm5505.

Pan, J.B., Hu, S.C., Shi, D., Cai, M.C., Li, Y.B., Zou, Q., and Ji, Z.L. (2013). PaGenBase: a pattern gene database for the global and dynamic understanding of gene function. PLoS One 8, e80747.

Parackova, Z., Zentsova, I., Bloomfield, M., Vrabcova, P., Smetanova, J., Klocperk, A., Meseznikov, G., Casas Mendez, L.F., Vymazal, T., and Sediva, A. (2020). Disharmonic Inflammatory Signatures in COVID-19: Augmented Neutrophils’ but Impaired Monocytes’ and Dendritic Cells’ Responsiveness. Cells 9.

Patel, S., Rauf, A., Khan, H., and Abu-Izneid, T. (2017). Renin-angiotensin-aldosterone (RAAS): The ubiquitous system for homeostasis and pathologies. Biomed Pharmacother 94, 317–325.

Pichlmair, A., Schulz, O., Tan, C.P., Naslund, T.I., Liljestrom, P., Weber, F., and Reis e Sousa, C. (2006). RIG-I-mediated antiviral responses to single-stranded RNA bearing 5’-phosphates. Science 314, 997–1001.

Pinero, J., Bravo, A., Queralt-Rosinach, N., Gutierrez-Sacristan, A., Deu-Pons, J., Centeno, E., Garcia-Garcia, J., Sanz, F., and Furlong, L.I. (2017). DisGeNET: a comprehensive platform integrating information on human disease-associated genes and variants. Nucleic Acids Res 45, D833–D839.

Ponti, G., Maccaferri, M., Ruini, C., Tomasi, A., and Ozben, T. (2020). Biomarkers associated with COVID-19 disease progression. Crit Rev Clin Lab Sci 57, 389–399.

Poulos, R.C., Hains, P.G., Shah, R., Lucas, N., Xavier, D., Manda, S.S., Anees, A., Koh, J.M.S., Mahboob, S., Wittman, M., et al. (2020). Strategies to enable large-scale proteomics for reproducible research. Nat Commun 11, 3793.

Qin, C., Zhou, L., Hu, Z., Zhang, S., Yang, S., Tao, Y., Xie, C., Ma, K., Shang, K., Wang, W., et al. (2020). Dysregulation of Immune Response in Patients With Coronavirus 2019 (COVID-19) in Wuhan, China. Clin Infect Dis 71, 762–768.

Raghunath, A., Sundarraj, K., Nagarajan, R., Arfuso, F., Bian, J., Kumar, A.P., Sethi, G., and Perumal, E. (2018). Antioxidant response elements: Discovery, classes, regulation and potential applications. Redox Biol 17, 297–314.

Raverdeau, M., and Mills, K.H. (2014). Modulation of T cell and innate immune responses by retinoic Acid. J Immunol 192, 2953–2958.

Rincon-Arevalo, H., Aue, A., Ritter, J., Szelinski, F., Khadzhynov, D., Zickler, D., Stefanski, L., Lino, A.C., Korper, S., Eckardt, K.U., et al. (2022). Altered increase in STAT1 expression and phosphorylation in severe COVID-19. Eur J Immunol 52, 138–148.

Riva, L., Yuan, S., Yin, X., Martin-Sancho, L., Matsunaga, N., Pache, L., Burgstaller-Muehlbacher, S., De Jesus, P.D., Teriete, P., Hull, M.V., et al. (2020). Discovery of SARS-CoV-2 antiviral drugs through large-scale compound repurposing. Nature 586, 113–119.

Ronit, A., Berg, R.M.G., Bay, J.T., Haugaard, A.K., Ahlstrom, M.G., Burgdorf, K.S., Ullum, H., Rorvig, S.B., Tjelle, K., Foss, N.B., et al. (2021). Compartmental immunophenotyping in COVID-19 ARDS: A case series. J Allergy Clin Immunol 147, 81–91.

Rooney, J., Oshida, K., Vasani, N., Vallanat, B., Ryan, N., Chorley, B.N., Wang, X., Bell, D.A., Wu, K.C., Aleksunes, L.M., et al. (2018). Activation of Nrf2 in the liver is associated with stress resistance mediated by suppression of the growth hormone-regulated STAT5b transcription factor. PLoS One 13, e0200004.

Rosenbaum, L. (2020). The Untold Toll - The Pandemic’s Effects on Patients without Covid-19. N Engl J Med 382, 2368–2371.

Rusinova, I., Forster, S., Yu, S., Kannan, A., Masse, M., Cumming, H., Chapman, R., and Hertzog, P.J. (2013). Interferome v2.0: an updated database of annotated interferon-regulated genes. Nucleic Acids Res 41, D1040–1046.

Rysz, S., Al-Saadi, J., Sjostrom, A., Farm, M., Campoccia Jalde, F., Platten, M., Eriksson, H., Klein, M., Vargas-Paris, R., Nyren, S., et al. (2021). COVID-19 pathophysiology may be driven by an imbalance in the renin-angiotensin-aldosterone system. Nat Commun 12, 2417.

Sanchez, J., Cohen, M., Zapater, J.L., and Eisenberg, Y. (2021). Primary Adrenal Insufficiency After COVID-19 Infection. AACE Clin Case Rep.

Sarohan, A.R., Akelma, H., Arac, E., Aslan, O., and Cen, O. (2022). Retinol Depletion in COVID-19. Clin Nutr Open Sci 43, 85–94.

Sarohan, A.R., Kizil, M., Inkaya, A.C., Mahmud, S., Akram, M., and Cen, O. (2021). A novel hypothesis for COVID-19 pathogenesis: Retinol depletion and retinoid signaling disorder. Cell Signal 87, 110121.

Schneider, W.M., Chevillotte, M.D., and Rice, C.M. (2014). Interferon-stimulated genes: a complex web of host defenses. Annu Rev Immunol 32, 513–545.

Schoggins, J.W., Wilson, S.J., Panis, M., Murphy, M.Y., Jones, C.T., Bieniasz, P., and Rice, C.M. (2011). A diverse range of gene products are effectors of the type I interferon antiviral response. Nature 472, 481–485.

Schulte-Schrepping, J., Reusch, N., Paclik, D., Bassler, K., Schlickeiser, S., Zhang, B.W., Kramer, B., Krammer, T., Brumhard, S., Bonaguro, L., et al. (2020). Severe COVID-19 Is Marked by a Dysregulated Myeloid Cell Compartment. Cell 182, 1419-+.

Shen, B., Yi, X., Sun, Y., Bi, X., Du, J., Zhang, C., Quan, S., Zhang, F., Sun, R., Qian, L., et al. (2020). Proteomic and Metabolomic Characterization of COVID-19 Patient Sera. Cell 182, 59–72 e15.

Solaimanzadeh, I. (2020). Acetazolamide, Nifedipine and Phosphodiesterase Inhibitors: Rationale for Their Utilization as Adjunctive Countermeasures in the Treatment of Coronavirus Disease 2019 (COVID-19). Cureus 12, e7343.

Sreeramkumar, V., Adrover, J.M., Ballesteros, I., Cuartero, M.I., Rossaint, J., Bilbao, I., Nacher, M., Pitaval, C., Radovanovic, I., Fukui, Y., et al. (2014). Neutrophils scan for activated platelets to initiate inflammation. Science 346, 1234–1238.

Takahashi, T., Ellingson, M.K., Wong, P., Israelow, B., Lucas, C., Klein, J., Silva, J., Mao, T., Oh, J.E., Tokuyama, M., et al. (2020). Sex differences in immune responses that underlie COVID-19 disease outcomes. Nature 588, 315–320.

Tan, M., Liu, Y., Zhou, R., Deng, X., Li, F., Liang, K., and Shi, Y. (2020). Immunopathological characteristics of coronavirus disease 2019 cases in Guangzhou, China. Immunology 160, 261–268.

Tay, M.Z., Poh, C.M., Renia, L., MacAry, P.A., and Ng, L.F.P. (2020). The trinity of COVID-19: immunity, inflammation and intervention. Nat Rev Immunol 20, 363–374.

Tepasse, P.R., Vollenberg, R., Fobker, M., Kabar, I., Schmidt, H., Meier, J.A., Nowacki, T., and Husing-Kabar, A. (2021). Vitamin A Plasma Levels in COVID-19 Patients: A Prospective Multicenter Study and Hypothesis. Nutrients 13.

Teuwen, L.A., Geldhof, V., Pasut, A., and Carmeliet, P. (2020). Author Correction: COVID-19: the vasculature unleashed. Nat Rev Immunol 20, 448.

Thimmulappa, R.K., Lee, H., Rangasamy, T., Reddy, S.P., Yamamoto, M., Kensler, T.W., and Biswal, S. (2006). Nrf2 is a critical regulator of the innate immune response and survival during experimental sepsis. J Clin Invest 116, 984–995.

Tong, L., Wang, L., Liao, S., Xiao, X., Qu, J., Wu, C., Zhu, Y., Tai, W., Huang, Y., Wang, P., et al. (2022). A Retinol Derivative Inhibits SARS-CoV-2 Infection by Interrupting Spike-Mediated Cellular Entry. mBio 13, e0148522.

Trasino, S.E. (2020). A role for retinoids in the treatment of COVID-19? Clin Exp Pharmacol Physiol 47, 1765–1767.

Ueland, T., Holter, J.C., Holten, A.R., Muller, K.E., Lind, A., Bekken, G.K., Dudman, S., Aukrust, P., Dyrhol-Riise, A.M., and Heggelund, L. (2020). Distinct and early increase in circulating MMP-9 in COVID-19 patients with respiratory failure. J Infect 81, e41–e43.

Vaduganathan, M., Vardeny, O., Michel, T., McMurray, J.J.V., Pfeffer, M.A., and Solomon, S.D. (2020). Renin-Angiotensin-Aldosterone System Inhibitors in Patients with Covid-19. N Engl J Med 382, 1653–1659.

Vanderboom, P.M., Mun, D.G., Madugundu, A.K., Mangalaparthi, K.K., Saraswat, M., Garapati, K., Chakraborty, R., Ebihara, H., Sun, J., and Pandey, A. (2021). Proteomic Signature of Host Response to SARS-CoV-2 Infection in the Nasopharynx. Mol Cell Proteomics 20, 100134.

Veras, F.P., Pontelli, M.C., Silva, C.M., Toller-Kawahisa, J.E., de Lima, M., Nascimento, D.C., Schneider, A.H., Caetite, D., Tavares, L.A., Paiva, I.M., et al. (2020). SARS-CoV-2-triggered neutrophil extracellular traps mediate COVID-19 pathology. J Exp Med 217.

Violi, F., Oliva, A., Cangemi, R., Ceccarelli, G., Pignatelli, P., Carnevale, R., Cammisotto, V., Lichtner, M., Alessandri, F., De Angelis, M., et al. (2020). Nox2 activation in Covid-19. Redox Biol 36, 101655.

Vollmy, F., van den Toorn, H., Zenezini Chiozzi, R., Zucchetti, O., Papi, A., Volta, C.A., Marracino, L., Vieceli Dalla Sega, F., Fortini, F., Demichev, V., et al. (2021). A serum proteome signature to predict mortality in severe COVID-19 patients. Life Sci Alliance 4.

Wang, S., Yao, X., Ma, S., Ping, Y., Fan, Y., Sun, S., He, Z., Shi, Y., Sun, L., Xiao, S., et al. (2021). A single-cell transcriptomic landscape of the lungs of patients with COVID-19. Nat Cell Biol 23, 1314–1328.

Webb, B.J., Peltan, I.D., Jensen, P., Hoda, D., Hunter, B., Silver, A., Starr, N., Buckel, W., Grisel, N., Hummel, E., et al. (2020). Clinical criteria for COVID-19-associated hyperinflammatory syndrome: a cohort study. Lancet Rheumatol 2, e754–e763.

Wilk, A.J., Rustagi, A., Zhao, N.Q., Roque, J., Martinez-Colon, G.J., McKechnie, J.L., Ivison, G.T., Ranganath, T., Vergara, R., Hollis, T., et al. (2020). A single-cell atlas of the peripheral immune response in patients with severe COVID-19. Nat Med 26, 1070–1076.

Wu, Z., Hu, R., Zhang, C., Ren, W., Yu, A., and Zhou, X. (2020). Elevation of plasma angiotensin II level is a potential pathogenesis for the critically ill COVID-19 patients. Crit Care 24, 290.

Xu, B., Lei, Y., Ren, X., Yin, F., Wu, W., Sun, Y., Wang, X., Sun, Q., Yang, X., Wang, X., et al. (2021). SOD1 is a Possible Predictor of COVID-19 Progression as Revealed by Plasma Proteomics. ACS Omega 6, 16826–16836.

Yang, D., Chu, H., Hou, Y., Chai, Y., Shuai, H., Lee, A.C., Zhang, X., Wang, Y., Hu, B., Huang, X., et al. (2020). Attenuated Interferon and Proinflammatory Response in SARS-CoV-2-Infected Human Dendritic Cells Is Associated With Viral Antagonism of STAT1 Phosphorylation. J Infect Dis 222, 734–745.

Ye, W., Chen, G., Li, X., Lan, X., Ji, C., Hou, M., Zhang, D., Zeng, G., Wang, Y., Xu, C., et al. (2020). Dynamic changes of D-dimer and neutrophil-lymphocyte count ratio as prognostic biomarkers in COVID-19. Respir Res 21, 169.

Zaid, Y., Dore, E., Dubuc, I., Archambault, A.S., Flamand, O., Laviolette, M., Flamand, N., Boilard, E., and Flamand, L. (2021). Chemokines and eicosanoids fuel the hyperinflammation within the lungs of patients with severe COVID-19. J Allergy Clin Immunol 148, 368–380 e363.

Zanoni, I. (2021). Interfering with SARS-CoV-2: are interferons friends or foes in COVID-19? Curr Opin Virol 50, 119–127.

Zhang, H., Penninger, J.M., Li, Y., Zhong, N., and Slutsky, A.S. (2020a). Angiotensin-converting enzyme 2 (ACE2) as a SARS-CoV-2 receptor: molecular mechanisms and potential therapeutic target. Intensive Care Med 46, 586–590.

Zhang, P., Zhu, L., Cai, J., Lei, F., Qin, J.J., Xie, J., Liu, Y.M., Zhao, Y.C., Huang, X., Lin, L., et al. (2020b). Association of Inpatient Use of Angiotensin-Converting Enzyme Inhibitors and Angiotensin II Receptor Blockers With Mortality Among Patients With Hypertension Hospitalized With COVID-19. Circ Res 126, 1671–1681.

Zhang, Q., Bastard, P., Liu, Z., Le Pen, J., Moncada-Velez, M., Chen, J., Ogishi, M., Sabli, I.K.D., Hodeib, S., Korol, C., et al. (2020c). Inborn errors of type I IFN immunity in patients with life-threatening COVID-19. Science 370.

Zhang, Z., Xing, X., Hensley, G., Chang, L.W., Liao, W., Abu-Amer, Y., and Sandell, L.J. (2010). Resistin induces expression of proinflammatory cytokines and chemokines in human articular chondrocytes via transcription and messenger RNA stabilization. Arthritis Rheum 62, 1993–2003.

Zheng, Y.Y., Ma, Y.T., Zhang, J.Y., and Xie, X. (2020). COVID-19 and the cardiovascular system. Nat Rev Cardiol 17, 259–260.

Zhou, Y., Zhou, B., Pache, L., Chang, M., Khodabakhshi, A.H., Tanaseichuk, O., Benner, C., and Chanda, S.K. (2019). Metascape provides a biologist-oriented resource for the analysis of systems-level datasets. Nat Commun 10, 1523.

