## Supplemental figures for "Proteome profiling of nasopharynx reveals pathophysiological signature of COVID-19 disease severity"

### Supplemental information

Supplemental information include four Figures and three Excel files. Each Excel file contains the table of information for patients' metadata and proteomics datasets respectively.

### Supplemental Figures

#### Figure S1 – DIA-based MS data acquisition approach

**A**) (**i**) Relative log expression of protein intensity after normalization and batch correction colored by equipment used. (**ii**) PCA plot of two principal components of protein intensities after normalization and batch correction colored by equipment used. (**iii**) Two-dimensional UMAP plot of protein intensities after normalization and batch correction colored by equipment used.

**B**) (**i**) Protein missing value matrix. (**ii**) Cumulative distribution of coefficient of variation calculated based on technical replicates.

**C**) (**i**) Number of identifications per minute of gradient time of single random sample. (**ii**) Scatterplot of retention time alignment between experimental values and library values. (**iii**) Scaled sum of intensities of individual samples in order of their acquisition indicating a single outlier of more than 10% change.

**D**) (**i**) Number of identifications per library fraction in each technical replicate. (**ii**) Number of proteins grouped by peptide count in the library. (**iii**) Number of proteins per protein group in the library.

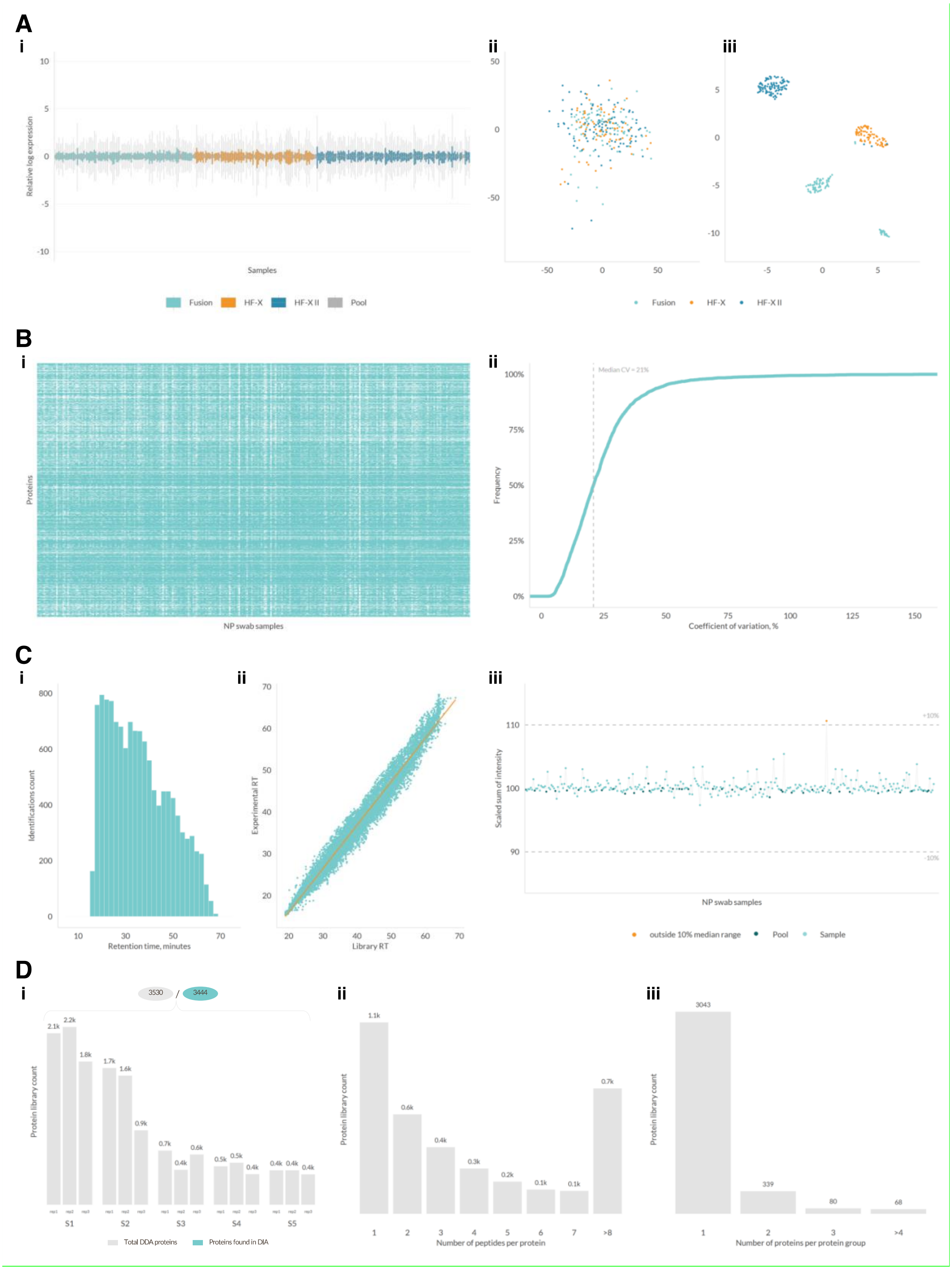
A
i
Relative log expression
Samples
Fusion
HF-X
HF-X II
Pool
ii
iii
Fusion
HF-X
HF-X II
B
i
Proteins
NP swab samples
ii
Median CV = 21%
Frequency
Coefficient of variation, %
C
i
Identifications count
Retention time, minutes
ii
Experimental RT
Library RT
iii
Scaled sum of intensity
+10%
-10%
NP swab samples
outside 10% median range
Pool
Sample
D
i
3530
3444
Protein library count
S1
S2
S3
S4
S5
Total DDA proteins
Proteins found in DIA
ii
Protein library count
Number of peptides per protein
iii
Protein library count
Number of proteins per protein group

**Figure S2 – Differential analysis by linear mixed effect modeling.**

**A**) Tissue or cell specific enrichment analysis of the list of differentially expressed proteins using PaGenBase database (Pan et al., 2013).

**B**) A circos plot representing a cross comparison between SA-M and SF-M differentially expressed protein lists, i.e. the overlap between the two protein lists only at the protein level, where purple curves link identical proteins.

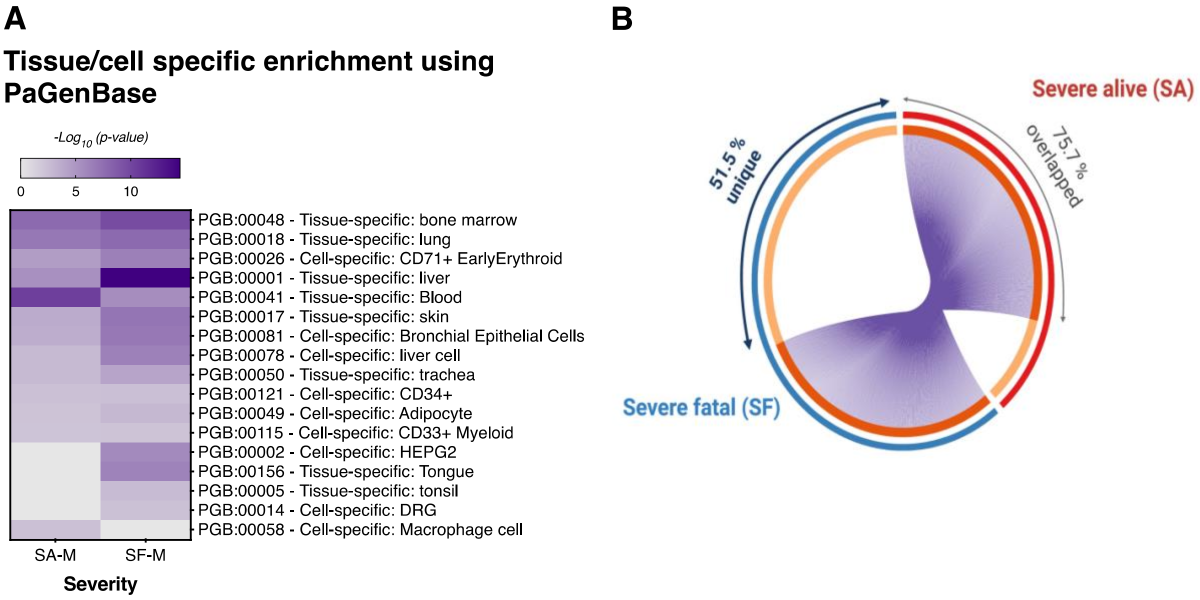
A
Tissue/cell specific enrichment using PaGenBase
-Log10 (p-value)
0
5
10
PGB:00048 - Tissue-specific: bone marrow
PGB:00018 - Tissue-specific: lung
PGB:00026 - Cell-specific: CD71+ EarlyErythroid
PGB:00001 - Tissue-specific: liver
PGB:00041 - Tissue-specific: Blood
PGB:00017 - Tissue-specific: skin
PGB:00081 - Cell-specific: Bronchial Epithelial Cells
PGB:00078 - Cell-specific: liver cell
PGB:00050 - Tissue-specific: trachea
PGB:00121 - Cell-specific: CD34+
PGB:00049 - Cell-specific: Adipocyte
PGB:00115 - Cell-specific: CD33+ Myeloid
PGB:00002 - Cell-specific: HEPG2
PGB:00156 - Tissue-specific: Tongue
PGB:00005 - Tissue-specific: tonsil
PGB:00014 - Cell-specific: DRG
PGB:00058 - Cell-specific: Macrophage cell
SA-M
SF-M
Severity
B
Severe alive (SA)
75.7 % overlapped
51.5 % unique
Severe fatal (SF)

**Figure S3 – Comparative analysis of selected differentially expressed proteins and their associated pathways in the nasopharynx of hospitalized COVID-19 patients.**

**A**) Protein network analysis of downregulated proteins involved in interferon signaling for SA-M and SF-M groups respectively.

**B - I**) Median-normalized protein abundance of downregulated NRF2 target proteins containing functional AREs (**B**), and upregulated proteins involved in neutrophil degranulation (**C**), interleukin signaling (**D**), platelet activation, signaling and aggregation (**E**), activation of matrix metalloproteinase enzymes (**F**), acute phase proteins (**G**), antioxidant proteins in ROS detoxification (**H**), and gaseous exchange of oxygen and carbon dioxide in erythrocytes (**I**) that are associated with COVID-19 disease severity (**i**) and gender (**ii**) respectively.

Data information: The *p*-values for violin plots (**B - I**) were calculated based on ordinary one-way ANOVA followed by Tukey’s multiple comparisons test for COVID-19 patients’ severity ($n_{M}$ = 92; $n_{SA}$ = 91; $n_{SF}$ = 90) and age groups ($n_{25\text{-}45}$ = 79; $n_{46\text{-}55}$ = 88; $n_{\geq 56}$ = 106), whereas the gender category ($n_{M}$ = 207; $n_{F}$ = 66) was analyzed using the unpaired t-test with Welch’s correction.

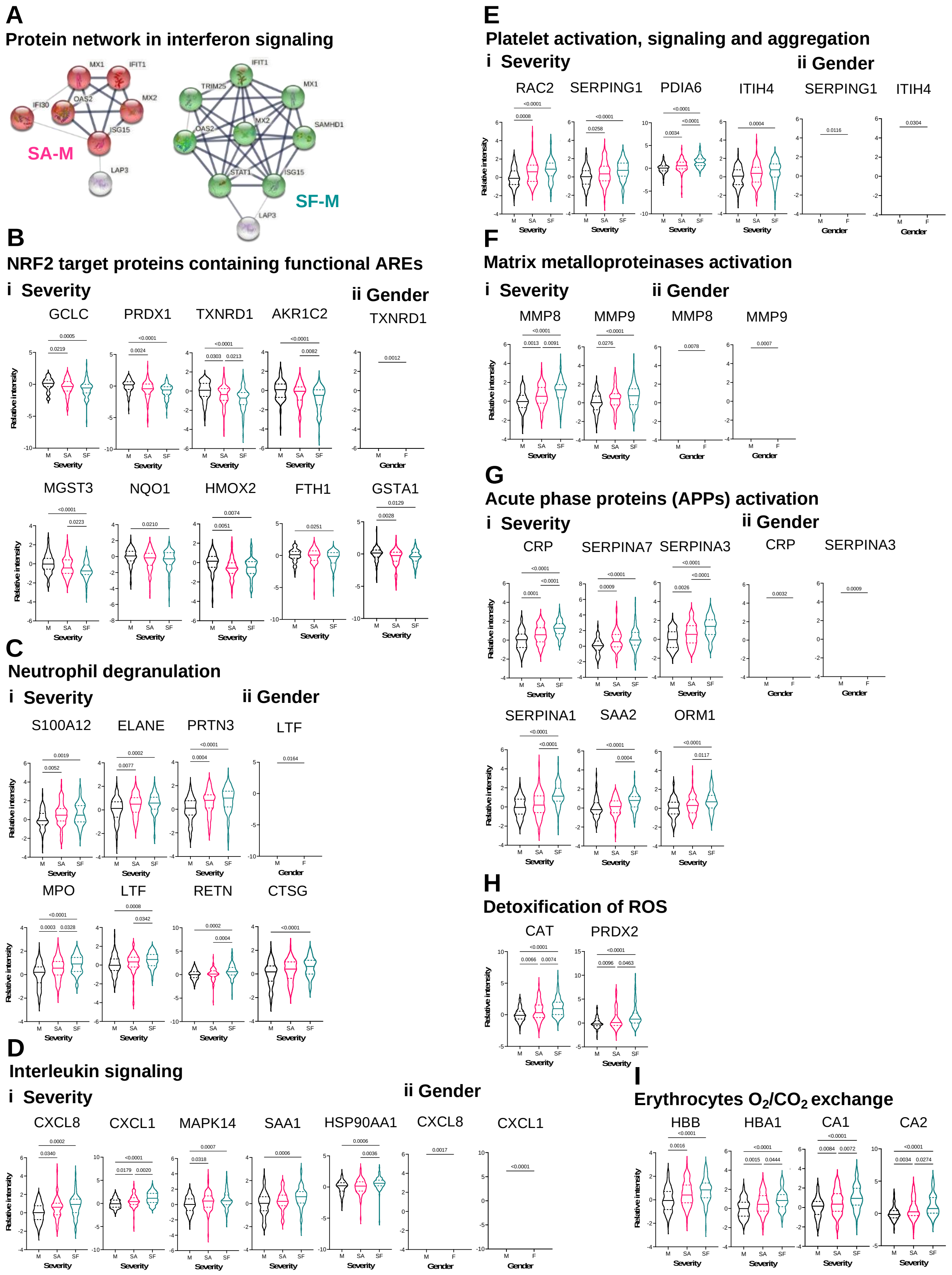
A
Protein network in interferon signaling
SA-M
SF-M
B
NRF2 target proteins containing functional AREs
i Severity
ii Gender
GCLC
PRDX1
TXNRD1
AKR1C2
TXNRD1
MGST3
NQO1
HMOX2
FTH1
GSTA1
C
Neutrophil degranulation
i Severity
ii Gender
S100A12
ELANE
PRTN3
LTF
MPO
LTF
RETN
CTSG
D
Interleukin signaling
i Severity
ii Gender
CXCL8
CXCL1
MAPK14
SAA1
HSP90AA1
CXCL8
CXCL1
E
Platelet activation, signaling and aggregation
i Severity
ii Gender
RAC2
SERPING1
PDIA6
ITIH4
SERPING1
ITIH4
F
Matrix metalloproteinases activation
i Severity
ii Gender
MMP8
MMP9
MMP8
MMP9
G
Acute phase proteins (APPs) activation
i Severity
ii Gender
CRP
SERPINA7
SERPINA3
CRP
SERPINA3
SERPINA1
SAA2
ORM1
H
Detoxification of ROS
CAT
PRDX2
I
Erythrocytes O2/CO2 exchange
HBB
HBA1
CA1
CA2
Relative intensity
Severity
Gender

**Figure S4 – Dysregulated proteins and their associated pathways in the nasopharynx of severe COVID-19 patients with fatal outcome.**

**A**) Protein function and pathway analysis of DE proteins unique to SF group itself. The (GO) enrichment statistical analysis was performed using Fisher's exact test with FDR multiple test correction, set at a critical value of FDR < 0.05. The number of proteins for each GO/Reactome term is annotated on the right side of the respective bar graphs.

**B**) (**i**) Enrichment analysis of all DE proteins for SA and SF using DisGeNET version 7.0. (**ii – iv**) Protein network analysis of selected DE proteins in SF group involved in antiviral defense, thrombocytopenia, vasculitis, interleukins signaling, neutrophil degranulation, inflammation, acute phase and bacterial infections using STRING version 11.5. The network nodes represent the proteins, while the line thickness indicate the degree of confidence prediction of functional association.

**C**) Median-normalized protein abundance of HP, PLG, FCER1G, ITGB2 and SERPINF1 involved in thrombocytopenia based on disease severity (**i**), gender (**ii**) and age (**iii**) groups respectively.

Data information: The *p*-values for violin plots were calculated based on ordinary one-way ANOVA followed by Tukey's multiple comparisons test for COVID-19 patients' severity ($n_{M}$ = 92; $n_{SA}$ = 91; $n_{SF}$ = 90) and age groups ($n_{25-45}$ = 79; $n_{46-55}$ = 88; $n_{\geq 56}$ = 106), whereas the gender category ($n_{M}$ = 207; $n_{F}$ = 66) was analyzed using the unpaired t-test with Welch's correction.

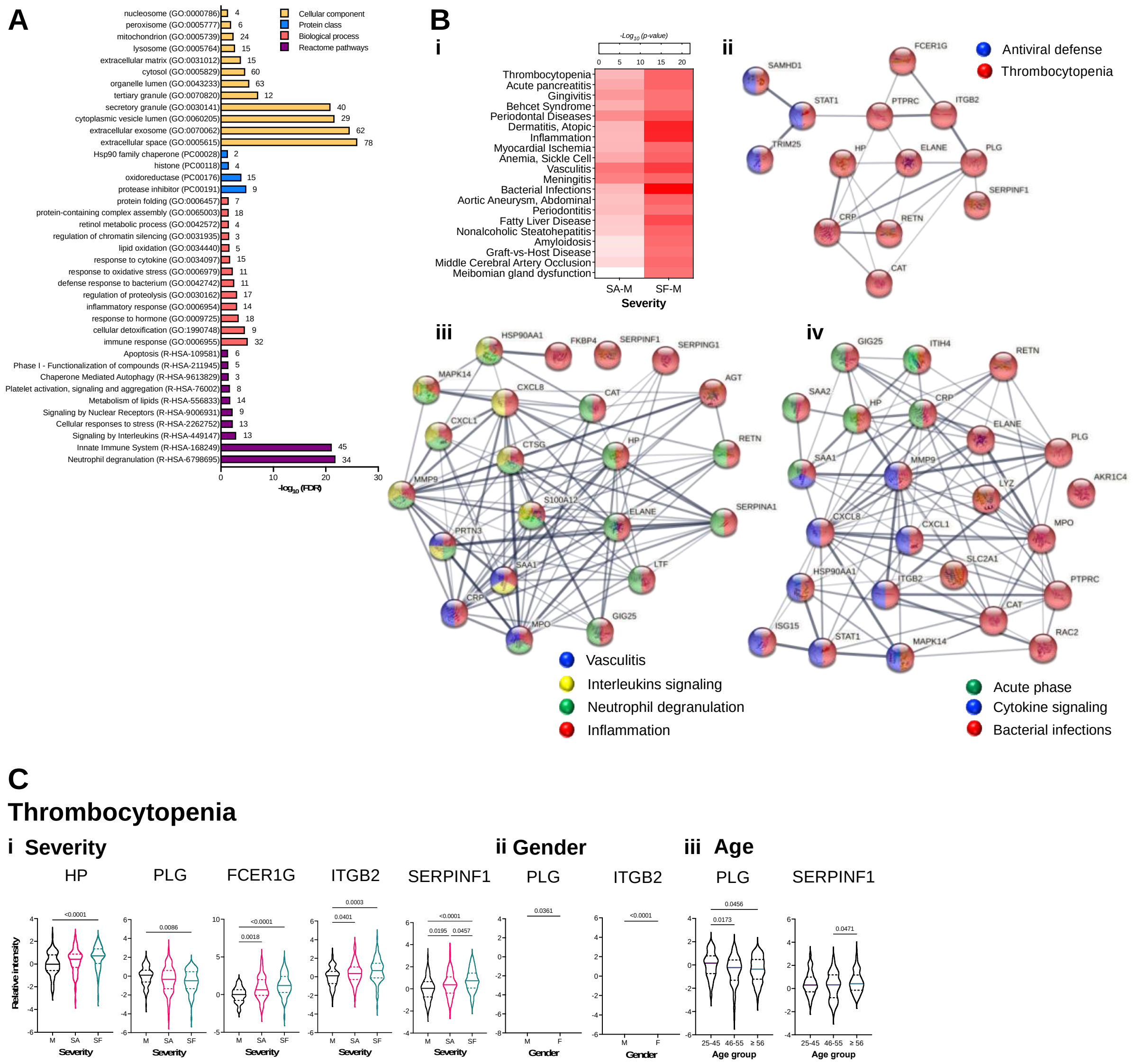
A
nucleosome (GO:0000786)
peroxisome (GO:0005777)
mitochondrion (GO:0005739)
lysosome (GO:0005764)
extracellular matrix (GO:0031012)
cytosol (GO:0005829)
organelle lumen (GO:0043233)
tertiary granule (GO:0070820)
secretory granule (GO:0030141)
cytoplasmic vesicle lumen (GO:0060205)
extracellular exosome (GO:0070062)
extracellular space (GO:0005615)
Hsp90 family chaperone (PC00028)
histone (PC00118)
oxidoreductase (PC00176)
protease inhibitor (PC00191)
protein folding (GO:0006457)
protein-containing complex assembly (GO:0065003)
retinol metabolic process (GO:0042572)
regulation of chromatin silencing (GO:0031935)
lipid oxidation (GO:0034440)
response to cytokine (GO:0034097)
response to oxidative stress (GO:0006979)
defense response to bacterium (GO:0042742)
regulation of proteolysis (GO:0030162)
inflammatory response (GO:0006954)
response to hormone (GO:0009725)
cellular detoxification (GO:1990748)
immune response (GO:0006955)
Apoptosis (R-HSA-109581)
Phase I - Functionalization of compounds (R-HSA-211945)
Chaperone Mediated Autophagy (R-HSA-9613829)
Platelet activation, signaling and aggregation (R-HSA-76002)
Metabolism of lipids (R-HSA-556833)
Signaling by Nuclear Receptors (R-HSA-9006931)
Cellular responses to stress (R-HSA-2262752)
Signaling by Interleukins (R-HSA-449147)
Innate Immune System (R-HSA-168249)
Neutrophil degranulation (R-HSA-6798695)
Cellular component
Protein class
Biological process
Reactome pathways
B
i
Thrombocytopenia
Acute pancreatitis
Gingivitis
Behcet Syndrome
Periodontal Diseases
Dermatitis, Atopic
Inflammation
Myocardial Ischemia
Anemia, Sickle Cell
Vasculitis
Meningitis
Bacterial Infections
Aortic Aneurysm, Abdominal
Periodontitis
Fatty Liver Disease
Nonalcoholic Steatohepatitis
Amyloidosis
Graft-vs-Host Disease
Middle Cerebral Artery Occlusion
Meibomian gland dysfunction
SA-M
SF-M
Severity
ii
Antiviral defense
Thrombocytopenia
iii
Vasculitis
Interleukins signaling
Neutrophil degranulation
Inflammation
iv
Acute phase
Cytokine signaling
Bacterial infections
C
Thrombocytopenia
i Severity
HP
PLG
FCER1G
ITGB2
SERPINF1
ii Gender
PLG
ITGB2
iii Age
PLG
SERPINF1
Relative intensity
Severity
Gender
Age group
